# A Liver-Targeted Copper Supplement Reduces Metabolic Dysfunction-Associated Liver Steatosis by Increasing Lipolysis and Fatty Acid Oxidation

**DOI:** 10.64898/2026.05.18.725917

**Authors:** Jaehee Kim, Vanha N. Pham, Timothy A. Su, Irene Liparulo, Diyala S. Shihadih, Tong Xiao, Xiao Xie, Yuichi Aki, Aidan T. Pezacki, Wendy Cao, James A. Olzmann, Joshua D. Rabinowitz, Andreas Stahl, Christopher J. Chang

## Abstract

Metabolic-associated steatotic liver disease (MASLD) is a prevalent liver disease driven by complex dysregulation of hepatic lipid metabolism. Here we show that copper deficiency is a nutrient vulnerability in steatotic liver disease and that selective liver-targeted copper supplementation can reduce excess lipid accumulation. Analysis of steatotic patient and mouse tissues identify widespread alterations in hepatic copper homeostasis markers. Integrated multi-omics analyses reveal that copper induces lipolysis of PLIN2-containing lipid droplets while lipid importer CD36 is downregulated. We show that copper inhibits cAMP hydrolase activity of PDE3B, thus activating PKA-mediated HSL and AMPK activation upstream of lipolysis. Fatty acids liberated through lipolysis are subsequently degraded via enhanced mitochondrial fatty acid oxidation, supported by energetic rewiring toward oxidative phosphorylation (OXPHOS) with increased copper-dependent complex IV and SOD1 activity. Our findings establish a multi-pronged mechanism by which hepatic copper supplementation coordinately regulates lipid metabolism in response to steatosis and unveils a therapeutic metallomedicine strategy to rewire lipid regulation.

**Summary:** Liver-targeted copper supplementation reduces diet-induced liver steatosis by dual activation of lipolysis and fatty acid degradation pathways.

**Graphical abstract:** 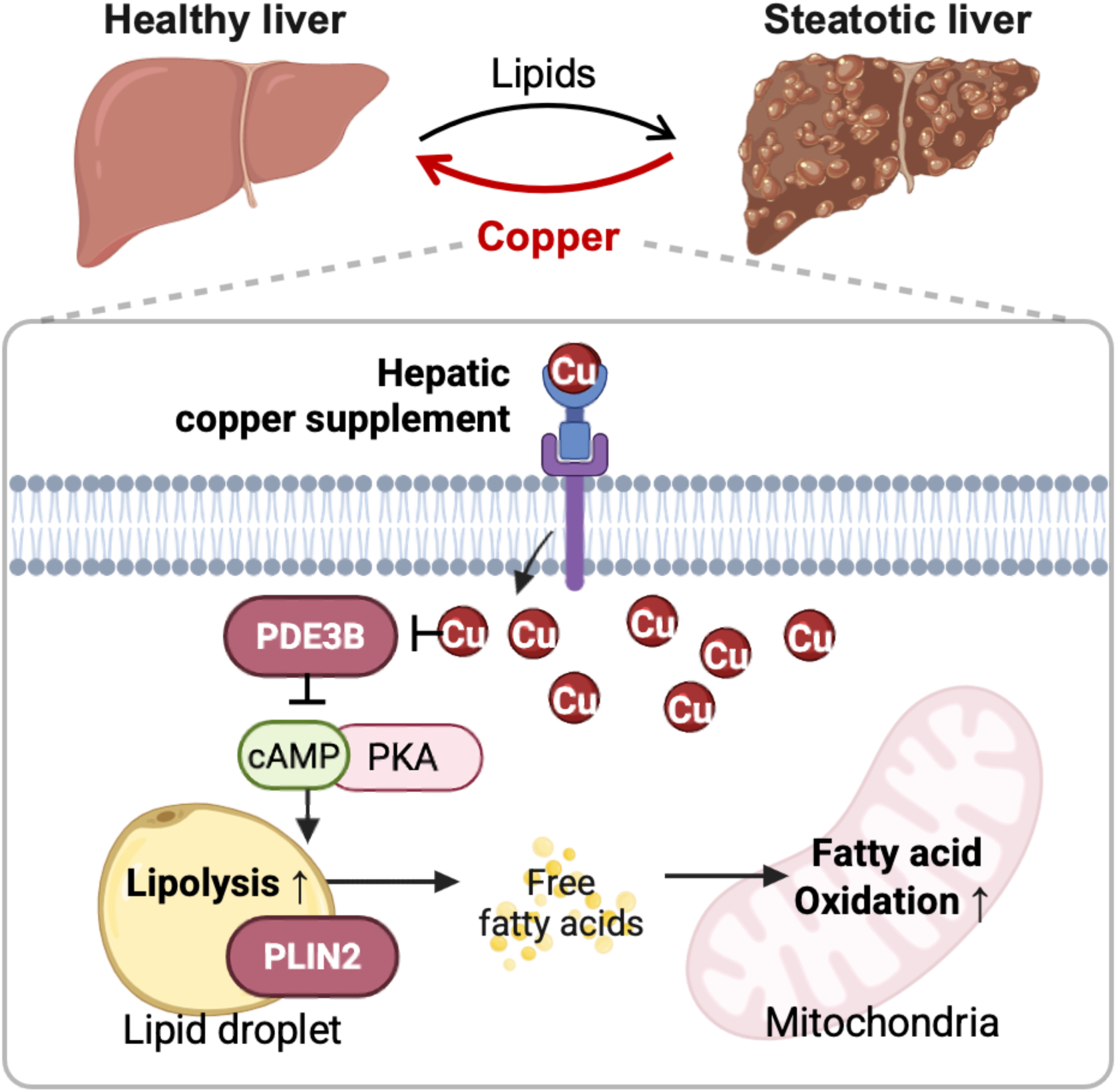

## Introduction

Metabolic-associated steatotic liver disease (MASLD) is a widespread human condition characterized by excess liver fat. As the most common liver disease in the world, MASLD affects up to 38% of all adults currently and is predicted to increase to more than 45% by 2050 (*1*). About 20% of MASLD patients develop metabolic-associated steatohepatitis (MASH), a more severe form of disease with higher risk of complications like fibrosis, cirrhosis, and liver cancer (1–3), along with increased likelihood of cardiovascular disorders (*4*). MASLD and MASH are complex, multilayered syndromes linked to metabolic dysfunction, such as in obesity and type 2 diabetes, with triggering factors including insulin resistance, oxidative stress, inflammation, and many others, making it difficult to decipher key biochemical mechanisms that underlie these diseases for therapeutic intervention (*3*, *5*).

In this context, copper is an essential metal nutrient acquired through the diet to maintain cellular and systemic health. The body requires tight regulation of copper import, export, and trafficking to sense and regulate a wide range of biological processes (*6–11*). Indeed, work from our laboratory and others have shown that copper can act not only as a static active site cofactor, but also as a dynamic regulator of cell signaling and metabolism, gene expression, and cell fate decisions by metalloallostery (*12*), which can give rise to distinctly characterized mechanisms of copper-dependent cell growth and cell death, termed cuproplasia (*13*) and cuproptosis (*14*), respectively (*15–21*). Along these lines, copper dysregulation mechanisms often occur in organ and cell-type specific contexts where copper overload or deficiency is connected to diseases, including genetic Wilson’s and Menkes diseases, cancer, neurodegenerative disorders, and cardiovascular failure (*22*). Interestingly, liver-localized copper deficiency has been reported in MASLD (*23–25*), but contributions of copper in the initiation and progression of steatotic liver diseases still remain elusive.

Here we show that liver copper deficiency is a key hallmark of steatotic metabolic liver diseases and a nutrient vulnerability that can be targeted for therapeutic intervention. This metal- and tissue-specific nutrient deficiency is accompanied by dysregulation of biochemical copper homeostasis machinery and an aberrant excess of liver fat. By selectively repleting liver copper with a hepatic-targeted small-molecule copper supplement, Gal-Cu(gtsm), we observed that diet-induced liver steatosis was effectively reduced to restore liver health. We showed that hepatic copper supplementation can regulate lipid droplet metabolism through metalloallosteric inhibition of phosphodiesterase 3B (PDE3B), which in turn increases cyclic AMP (cAMP) and protein kinase A (PKA) signaling to both enhance lipolysis and suppress lipid import. Liver copper repletion in dietary models of MASLD also reprogrammed metabolism toward mitochondrial respiration and fatty acid oxidation to further process lipid degradation, with concomitant increases in antioxidant activity of copper-dependent superoxide dismutase 1 (SOD1). Taken together, our work identifies copper as a nutrient vulnerability in fatty liver disease and unveils tissue-specific metal supplementation a therapeutic strategy that can be uniquely leveraged to regulate lipid catabolism across multiple pathways spanning lipolysis, lipid uptake, and fatty acid oxidation to reduce steatosis.

## Results

### Hepatic copper is depleted with dysregulation of copper homeostasis during MASLD and MASH progression *in vivo*

To build on observations of copper deficiency in fatty liver patients (*23*), we examined human RNA-seq datasets from liver biopsies (MASLD = 313, MASH = 539, CTRL = 109, total = 1050 samples) (*26*) and identified genetic signatures of altered copper homeostasis in steatotic liver disease patients (Fig. 1A and Table S1). Gene ontology (GO) analysis revealed that human MASLD livers show significant alteration of cation transmembrane transporter activity, dicarboxylic acid transporter activity, and ATPase activator activity, along with expected changes in glucan metabolic process and insulin-like growth receptor binding pathways (Fig. 1B). Interestingly, serine/threonine kinase activity, oxygen binding, and proton-transporting ATPase activity genes were significantly altered as well. The more progressed MASH livers showed alterations of genes in response to metal ion, dicarboxylic acid metabolism, and cellular response to copper ion pathways, as well as symptom-relevant pathways such as regulation of cytokinesis, collagen fibril organization, and complement activation (Fig. 1C). Unlike MASLD livers, MASH livers also showed significant changes in aspartate family metabolic processes and L-glutamate transmembrane transport, as well as vitamin B6 binding pathways. These pathways support the connection of metal ion status, particularly of copper, to liver disease.

**Fig. 1:**
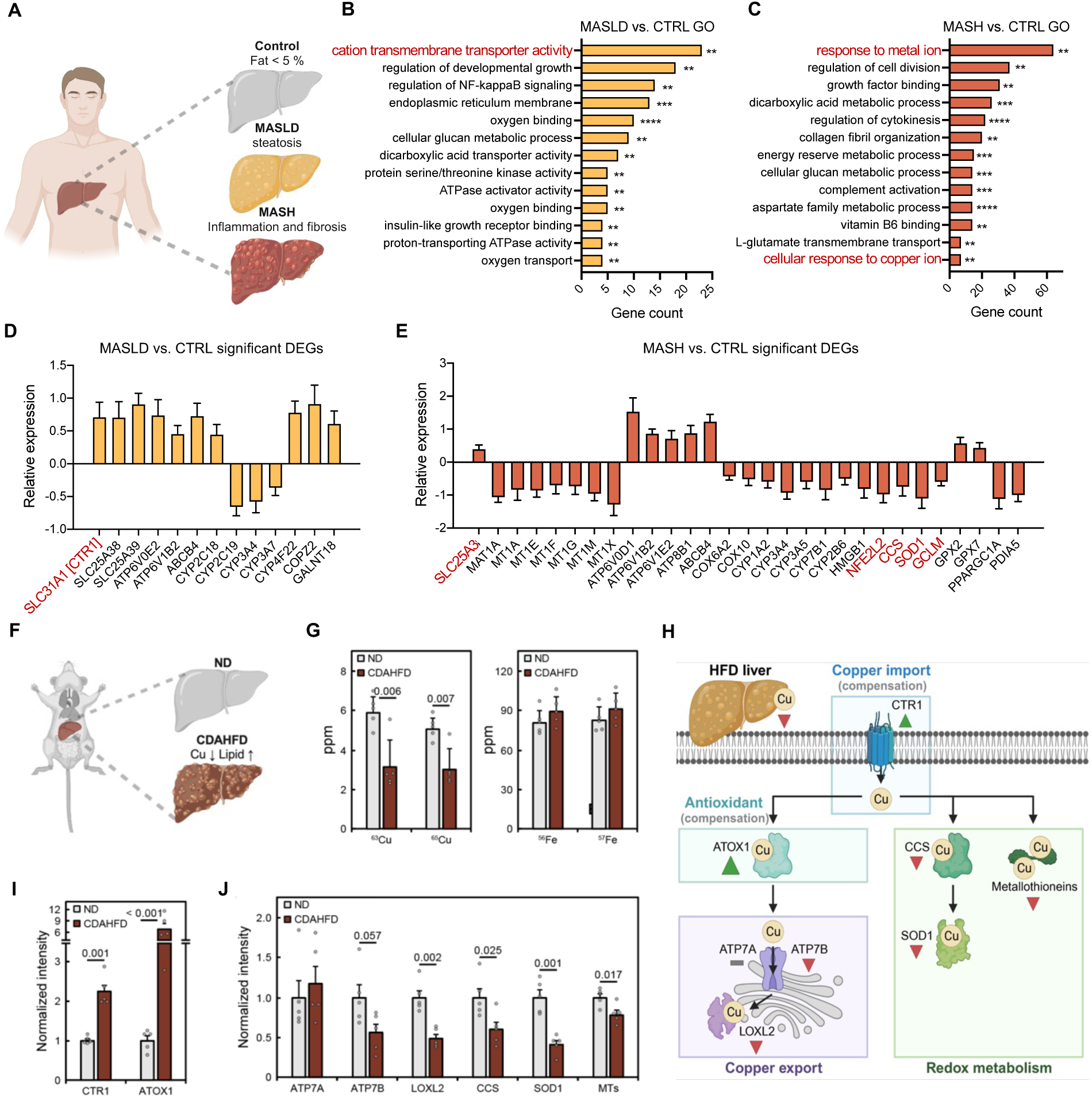
Hepatic copper is depleted during MASLD and MASH progression *in vivo*. **A.** Overview of metabolic liver disease progression. Hepatic lipid accumulation is highlighted in MASLD and MASH progression while copper gets depleted. **B and C.** Gene ontology (GO) analysis results in MASLD and MASH compared to healthy individuals. Red highlights pathways related to copper. Unpaired t-test; ns, not significant, ∗p < 0.05, ∗∗p < 0.01, ∗∗∗p < 0.001, ∗∗∗∗p < 0.0001. **D and E.** Copper-mediated genes with significant fold changes (one-way Anova, p-value < 0.05) in MASLD and MASH compared to healthy individuals. All error bars represent mean ± SD and red highlights genes related to copper homeostasis. **F.** Schematic of copper and lipid alteration comparing normal diet (ND) and choline-deficient, L-amino acid-defined, high-fat diet (CDAHFD). **G.** Hepatic metal levels measured by ICP-MS in CDAHFD mouse livers. **H.** Overview of copper homeostasis markers in various pathways. **I and J.** Copper homeostasis markers whose expression changed in CDAHFD livers.

Next, we investigated individual differentially expressed genes that showed statistical significance (adjusted p-value < 0.05, Fig 1D and E). Human MASLD and MASH livers showed significant increases in the plasma membrane copper uptake protein SLC31A1 (CTR1) and the mitochondrial copper and phosphate transporter SLC25A3 compared to healthy livers, while metallothionein genes were downregulated in MASH, likely reflecting limited copper availability. Various genes in ATPase family were upregulated in both hMASLD and hMASH, but expression levels of copper-dependent ATPases ATP7A or ATP7B did not appear to be significantly changed. However, ABCB4 and ABCB1 significantly increased in MASLD and MASH, respectively, showing cellular stress responses caused by copper imbalances in these patient livers (15, 16). Moreover, CYP family genes, which are known to be activated by copper-induced stress responses (*29*), significantly decreased in both MASLD and MASH livers. Furthermore, well-studied copper-binding proteins such as the antioxidant protein superoxide dismutase 1 (SOD1) and its metallochaperone CCS (*30*), as well as genes involved in biosynthesis of glutathione (GSH), which serves to buffer intracellular labile copper pools (*31*), such as NFE2L2 and GCLM (*32*), were significantly downregulated in hMASH livers. Interestingly, SLC25A39, GPX2, and GPX7 increased, possibly to compensate the downregulation of antioxidants SOD1 and GSH in hMASH livers. Taken together, analysis of human MASLD and MASH tissues establish that copper homeostasis is altered in steatotic liver diseases.

To further study how copper is dysregulated in metabolic liver diseases, we analyzed mice livers from a choline-deficient, L-amino acid-defined, high-fat diet (CDAHFD), which models the symptoms of severe fibrosis and inflammation of MASH (Fig. 1F) (*33*, *34*). Hepatic copper levels were measured by laser ablation ICP-MS, showing a 1.8-fold decrease in copper in the CDAHFD samples compared to normal diet (ND) (Fig. 1G), with no significant change in iron levels. Next, a panel of copper homeostasis proteins were compared by immunoblotting (Fig. 1H-J and Fig. S1A). The expression levels of the copper uptake protein CTR1, as well as the copper metallochaperone and antioxidant protein ATOX1, were significantly increased in CDAHFD livers compared to ND livers, which suggest a compensatory response to copper deficiency in CDAHFD livers (Fig. 1I). In contrast, the copper exporter protein ATP7B, copper metallochaperone for superoxide dismutase (CCS), and metallothioneins (MTs) that store copper were all downregulated in CDAHFD livers relative to ND controls, consistent with a decrease in copper bioavailability. Moreover, the expression levels of copper-dependent enzymes lysyl oxidase like-protein 2 (LOXL2) and SOD1, as well as the SOD1 enzyme activity, were significantly decreased in CDAHFD compared to control ND livers (Fig. 1J and Fig. S1B). Interestingly, expression changes in these copper homeostasis proteins were more pronounced than classic lipid metabolism proteins such as DGAT2, FABP1, ATGL1, ABHD5, or PNPLA4 in HFD livers (Table. S1). These mouse data are aligned with the human patient analysis and show altered liver copper homeostasis and protein expression changes to compensate for copper deficiency.

### Liver-targeted copper supplementation repletes hepatic copper deficiency and prevents diet-induced steatosis

Given observations of liver-localized copper deficiency and associated alterations in copper homeostasis protein expression across human and mouse MASLD and MASH models, we reasoned that targeted copper supplementation to liver tissue might have beneficial effects to ameliorate lipid dysregulation in these diseases. Coupled with previous observations from our laboratory using a caged luciferin probe for activity-based sensing of copper that showed a liver-localized copper deficiency in a diet-induced mouse model of MASLD in as early as two weeks (*24*), before substantial weight gain is observed, we sought to evaluate liver-directed copper supplementation as a prophylactic MASLD treatment strategy.

As a starting point for testing this approach, our laboratory previously reported a small-molecule compound for liver-specific copper delivery, Gal-Cu(gtsm), by linking a Cu(gtsm) copper ionophore fragment to an *N*-acetylgalactosamine (GalNAc) warhead to target ASGPR (*35*). The hepatic-targeted copper ionophore is designed to bypass the ubiquitously expressed copper importer CTR1, enabling selective copper accumulation in the liver upon ASGPR-mediated ionophore uptake and lysosome-mediated copper release (Fig. 2A). The receptor-dependent delivery of copper in this manner minimizes oxidative stress that can result from non-targeted systemic copper supplementation. Along these lines, analysis of hMASLD data showed upregulation of GALNT18, which increases endocytosis mediated by the liver-specific asialoglycoprotein receptor (ASGPR), which benefits this therapeutic strategy to MASLD mouse model (Fig. 1D). We first tested the toxicity of Gal-Cu(gtsm) compared to Cu(gtsm) to confirm the phenotype of regulated copper release. As expected, regulated copper delivery to Huh7 cells using Gal-Cu(gtsm) exhibited no significant increase of cytoplasmic or mitochondrial reactive oxygen species (Fig. S1D and E). In contrast, the non-targeted Cu(gtsm) ionophore triggered an increase in superoxide, leading to heightened cytotoxicity as measured by live/dead-cell imaging. Moreover, treatment of mice *in vivo* with Gal-Cu(gtsm) twice per week for 10 weeks by intraperitoneal injection showed no noticeable toxicity indicated by any gross morphological abnormalities at a dose of 7.26 mg Gal-Cu(gtsm)/kg mouse twice per week, which is equivalent to 0.75 mg Cu/kg mouse per dose (Fig. S1F).

**Fig. 2:**
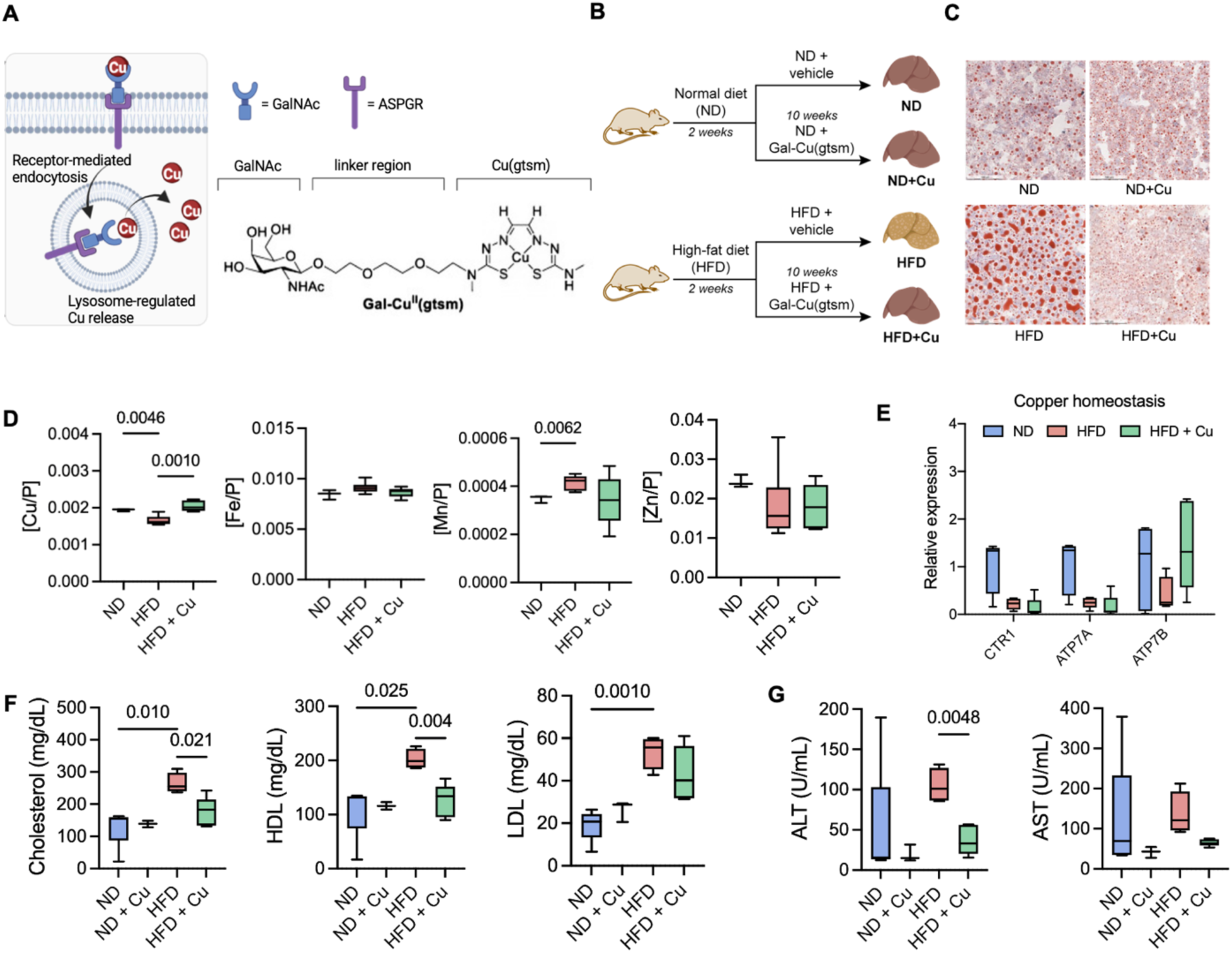
Gal-Cu(gtsm) selectively repletes hepatic copper deficiency and prevents diet-induced steatosis. **A.** Gal-Cu(gtsm) structure and mechanism of action for liver-targeted copper delivery. **B.** Overview of the prophylactic study of hepatic copper repletion by Gal-Cu(gtsm) in normal diet (ND) and high-fat diet (HFD) steatosis model with copper supplementation (+Cu) (ND, HFD+Cu, n = 6; ND+Cu, HFD, n = 5). **C.** Oil red O staining of liver sections at the endpoint of the feeding experiments. **D.** Metal measurements in livers normalized to phosphorus using ICP-MS. **E.** Relative protein expression of key copper homeostasis markers whose expression significantly changed in livers. **F.** Cholesterol and lipoproteins measured in serum. **G.** Liver enzyme activities measured in serum. All error bars represent mean ± SD. Unpaired t-test; ns, not significant, ∗p < 0.05, ∗∗p < 0.01, ∗∗∗p < 0.001, ∗∗∗∗p < 0.0001.

With these data in hand, we moved to evaluate the use of Gal-Cu(gtsm) for prophylactic MASLD treatment. Mice were fed with a normal diet (ND) or high-fat diet (HFD) for 2 weeks, at a time point at which the HFD cohorts show liver copper deficiency, followed by 10 weeks with or without Gal-Cu(gtsm) treatment (Fig. 2B). We were pleased to observe that liver-targeted copper supplementation prevented buildup of excess fat in livers of the HFD mouse models treated with Gal-Cu(gtsm) relative to untreated HFD controls (Fig. 2C). As expected, hepatic copper levels from HFD group were decreased compared to ND controls, and Gal-Cu(gtsm) treatment rescued this copper deficiency, with no significant changes in other transition metal nutrients (Fig. 2D). In agreement with previous findings (Fig. 1J), both ATP7A and ATP7B were significantly decreased in HFD mice compared to ND controls, whereas Gal-Cu(gtsm) copper supplementation restored ATP7B expression back to normal levels (Fig. 2E). Notably, Gal-Cu(gtsm)-treated mice exhibited reduced total blood cholesterol and low-density lipoprotein (LDL) cholesterol levels, without changes in high-density lipoprotein (HDL) cholesterol, indicating diminished hepatic cholesterol export while preserving systemic cholesterol clearance capacity upon restoration of hepatic copper levels (Fig. 2F). Alongside improved lipid regulation, markers of liver health, as assessed by aspartate transaminase (AST) and alanine transaminase (ALT) enzymatic activity, were significantly improved following copper supplementation (Fig. 2G), demonstrating that hepatic copper repletion mitigates steatosis-associated pathology and disease progression.

### Liver-targeted copper supplementation rescues diet-induced steatosis

We next examined whether hepatic copper repletion could rescue diet-induced steatosis. To this end, mice were fed with a high-fat diet (HFD) for 12 weeks, followed by 8 weeks of normal diet (ND) with or without Gal-Cu(gtsm) treatment (Fig. 3A). We were pleased to observe that even after full progression of steatosis after HFD feeding, hepatic-directed copper supplementation was able to reduce excess fat accumulation in the liver compared to untreated controls (Fig. 3B and Fig. S2A). In addition, during the course of Gal-Cu(gtsm) supplementation we noted a decrease in weight gain compared to the HFD control group (Fig. 3C, D, and Fig. S2B). Importantly, EchoMRI revealed that the improvement in weight control in HFD+Cu group was attributable mainly from mice gaining less fat mass, rather than compromising lean mass relative to the HFD group (Fig. 3E). Moreover, hepatic copper repletion after full progression of steatosis lowered blood glucose levels (Fig. 3F) without decreasing food consumption (Fig. 3G), reduced the wet mass of livers, and restored activities of AST and ALT liver enzymes back down to normal levels (Fig. 3H and I).

**Fig. 3:**
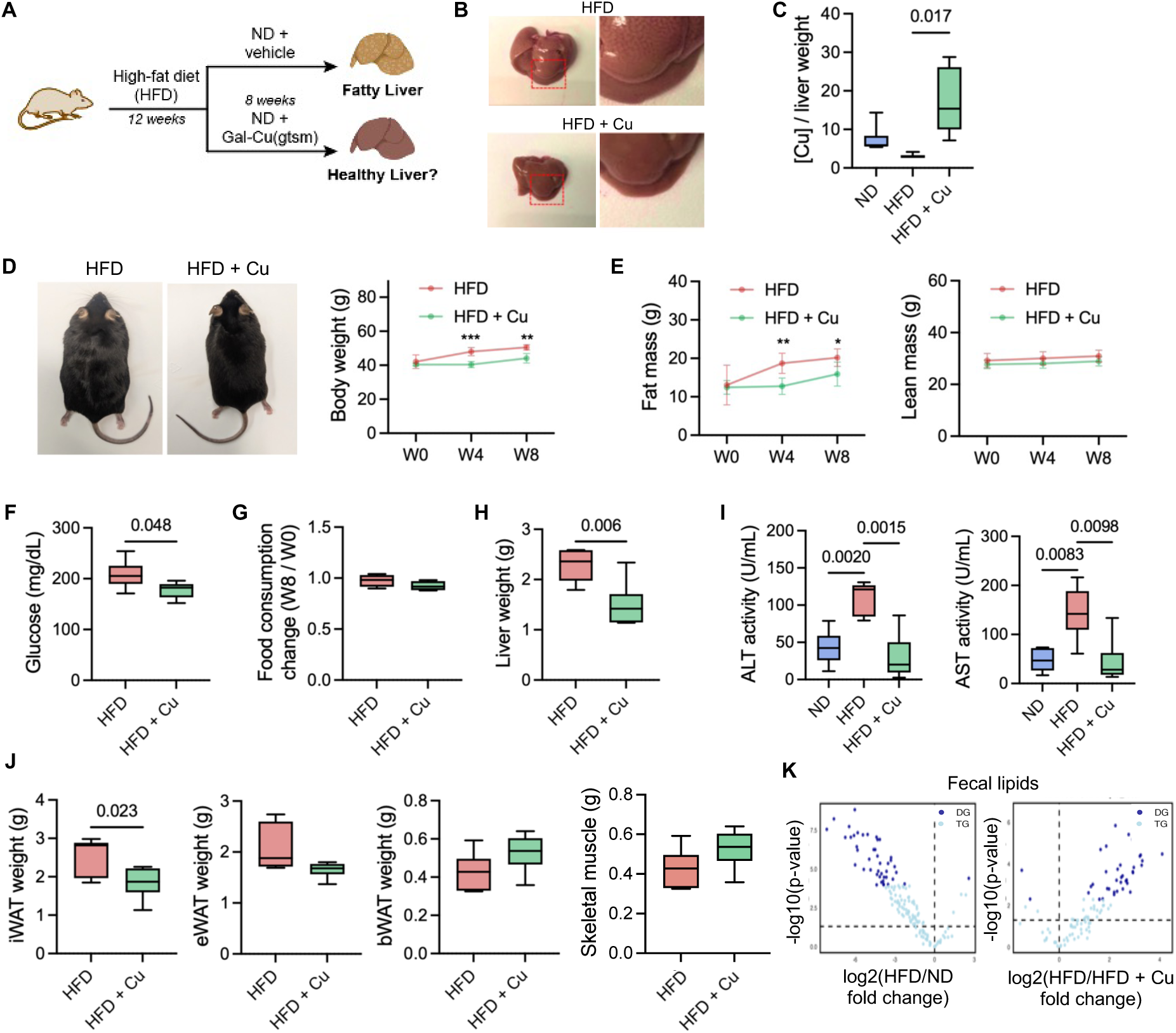
Liver-targeted copper supplementation rescues diet-induced steatosis and restores liver health. **A.** Overview of the therapeutic study of hepatic copper repletion by Gal-Cu(gtsm) in steatosis (HFD) dietary models (n = 6 per group). **B.** Representative pictures of wet livers dissected from mice in the study with or without visible lipid accumulation. **C.** ICP-MS measurements of copper at the study endpoint. **D.** Representative pictures of mice at the endpoint of the experiment and body weights measured at the start of Gal-Cu(gtsm) treatment (W0) and the eight weeks of treatment (W8). **E.** EchoMRI measurements of fat and lean mass at 0, 4, and 8 weeks of Gal-Cu(gtsm) treatment. **F.** Serum glucose levels measured at week 8 of Gal-Cu(gtsm) or DPBS treatment. **G.** Food consumption change measured by cage at week 0 and 8 of Gal-Cu(gtsm) or DPBS treatment. **H.** Wet liver mass measured after sacrifice (n=6 per group). **I.** Serum ALT and AST activities measured (n=6 per group). **J.** Wet tissue mass of white and brown adipose tissue. **K.** Volcano plots of diglycerides (DG) and triglycerides (TG) measurements in fecal pellets from mice fed with ND and HFD collected after 48 hours after copper supplementation on week 8. All error bars represent mean ± SD. Unpaired t-test; ns, not significant, ∗p < 0.05, ∗∗p < 0.01, ∗∗∗p < 0.001, ∗∗∗∗p < 0.0001.

Hepatic copper supplementation also decreased the wet mass of iWAT (inguinal white adipose tissue) and eWAT (epididymal white adipose tissue) relative to HFD group. While unexpectedly, BAT (brown adipose tissue) and skeletal muscle slightly increased suggesting that copper supplementation does not compromise mice health to reduce weight but rather improves it (Fig. 3J and Fig. S2C). To better understand these phenomena, lipidomics PCA revealed a noticeable shift in the liver lipidome of the copper-supplemented HFD group compared with the HFD controls, while the plasma lipidome mostly overlapped, suggesting that copper-induced lipid profile alterations occur mainly within liver and do not significantly affect the composition of lipids exported into plasma (Fig. S2D and E). Moreover, the lipid profiles in iWAT, eWAT and BAT were largely unaffected by copper supplementation despite changes in their wet masses, suggesting that hepatic-targeted copper repletion does not directly alter lipid compositions in tissues beyond liver but may benefit regulating lipid circulation and storage in neighboring adipose tissues (Fig. S2F). We then examined how copper supplementation alters lipid excretion by performing lipidomics analyses on fecal pellets. Again, mice were fed with ND, HFD or HFD + Cu and fecal pellets were collected after 48 hours. Notably, the excretion of TG and DG fatty acids decreased in HFD mice compared to ND to explain increased lipid uptake and storage, while this excretion further dropped when copper was supplemented ruling out the possibility that copper supplementation regulates fat gain by increased lipid excretion (Fig. 3K and Fig. S2G). The collective data establishes that hepatic copper supplementation selectively alters hepatic lipid regulation relative to other tissues to promotes resolution for diet-induced steatosis during diet normalization.

### Multi-omics profiling characterizes biochemical signatures of aberrant liver lipid metabolism rescued by hepatic copper repletion

To gain further biochemical insights into how hepatic-targeted copper supplementation reduces liver steatosis, we performed systematic, multi-omics analyses at the proteome, metabolome, and lipidome level in mouse MASLD models. First, we looked at 12 MASLD-associated genes, including the antioxidant response gene NFE2L2 (NRF2), to determine whether copper regulates lipids at a genetic level. However, RT-qPCR analyses showed little to no consistent patterns in these MASLD-associated transcripts between the copper-supplemented and untreated control groups, suggesting that copper does not directly alter lipid metabolism at the transcriptional level (Fig. S3 and Table S2).

We next profiled the liver proteomes, where principal component analysis (PCA) showed shifts across groups (Fig. 4A and Table S3). Notably, Perilipin 2 (PLIN2), a protein involved in lipid droplet formation and function (*36*, *37*); CD36, a major fatty acid transporter facilitating the uptake of long-chain fatty acids into cells (*38*); and the acyl-CoA thioesterase 1 (ACOT1) that breaks down acyl-CoA into free fatty acids (*39*, *40*) were all significantly upregulated by HFD relative to ND and reduced back down to normal levels by hepatic copper supplementation (Fig. 4B and Fig. S4). On the other hand, Fatty Acid-Binding Protein 5 (FABP5), a master lipid chaperone that carries long-chain fatty acids to lipid droplets to regulate lipolysis (*41*, *42*), was also significantly downregulated by HFD compared to ND but not affected by Gal-Cu(gtsm) (Fig. S5). In addition, lipid detoxification regulators, including GSTM and GSTA families (*43*) and the ferroptosis suppressor GPX4 (*44*, *45*), were increased by HFD and decreased by Gal-Cu(gtsm) treatment, which can be attributed to cellular responses related to GSH production. Indeed, REACTOME enrichment analysis revealed that copper-supplemented livers show altered expressions of lipid metabolism, fatty acid oxidation, and biological oxidation (Fig. 4C).

**Fig. 4:**
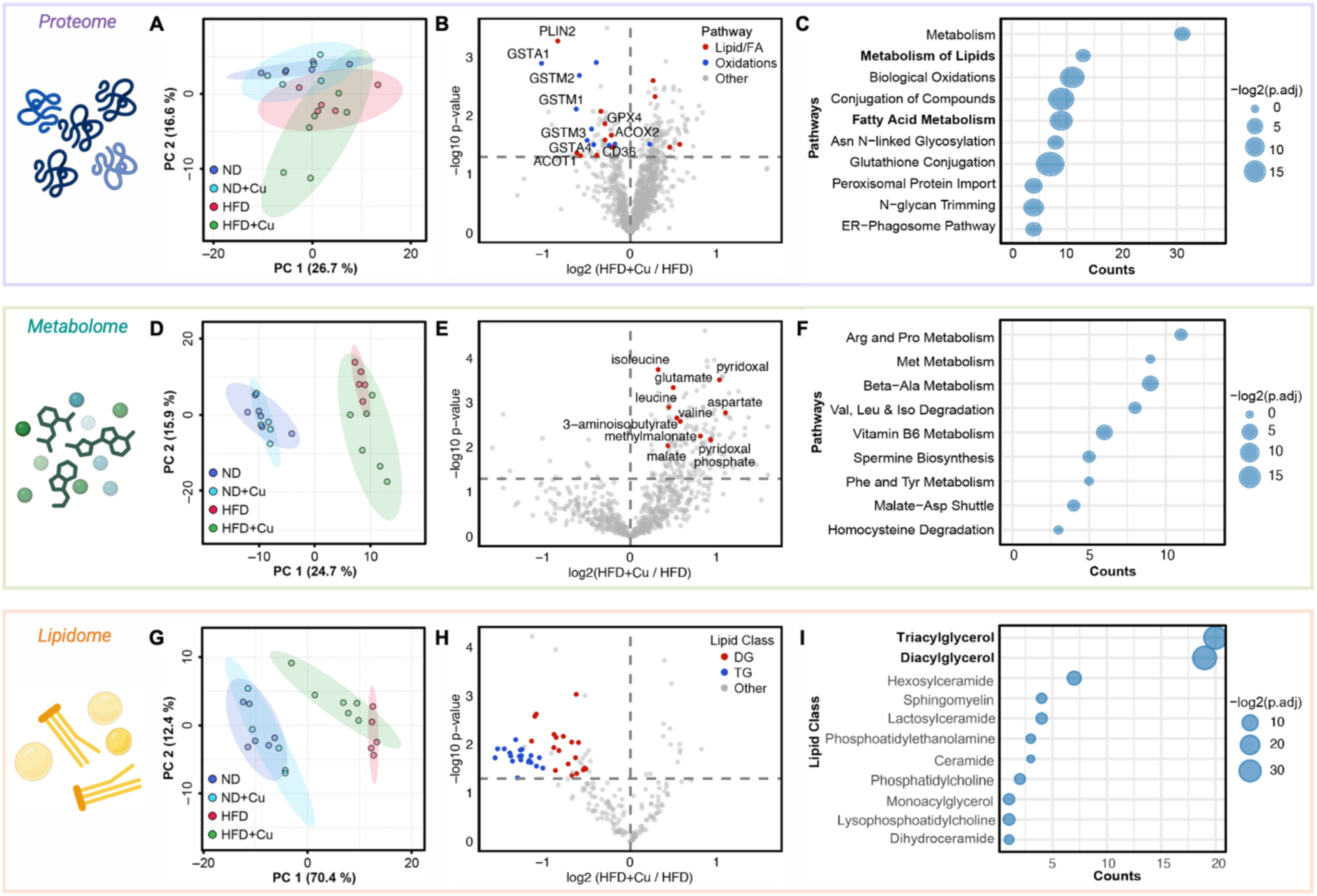
Multi-omics profiling characterizes biochemical signatures of aberrant liver lipid metabolism rescued by hepatic copper supplementation. **A-I.** Analysis of liver from mice that were fed with ND or HFD for 2 weeks followed by treatment with Gal-Cu(gtsm) or DPBS vehicle for the following 10 weeks (Fig. 2B, ND, HFD+Cu, n = 6; ND+Cu, HFD, n = 5 per group). **A.** PCA of mouse liver proteomics. **B.** Volcano plot of mouse liver proteomics comparing HFD+Cu to HFD. Points from Metabolism of Lipids and Fatty Acid Metabolism (Lipid/FA) and Biological Oxidations (Oxidations) are colored with mentioned proteins labeled. **C.** Pathway analysis of differentially expressed proteins in HFD+Cu compared to HFD revealed lipid metabolism and fatty acid metabolism as enriched pathways. **D.** PCA of mouse liver metabolomics. **E.** Volcano plot of mouse liver metabolomics comparing HFD+Cu to HFD. Points from Val, Leu, and Iso Degradation and Malate-Asp Shuttle are colored and labeled. **F.** Pathway analysis of differentially expressed metabolites in HFD+Cu compared to HFD. **G.** PCA of mouse liver lipidomics. **H.** Volcano plot of mouse liver lipidomics comparing HFD+Cu to HFD shows generally reduced DG and TG by copper supplementation. Points are colored by select lipid classes. **I.** Lipid class enrichment analysis of differentially expressed lipids in HFD+Cu compared to HFD.

We then profiled the liver metabolomes (Fig. 4D, Table S4 and Fig. S6 and S7). Key metabolites that showed a decrease in HFD compared to ND but restored back to normal levels by copper supplementation include cAMP, choline, glutamate, and retinol (vitamin A). Importantly, Gal-Cu(gtsm) treatment increased retinol, which can help transport cholesterol (*46*, *47*), as well as choline, an essential precursor for the biosynthesis of phosphatidylcholine (PC), which aids in lipid export from the liver and whose deficiency is a key hallmark of MASH (*48–50*). Moreover, glutathione was also decreased in HFD and restored by copper, again implying that this metal nutrient supplementation increases GSH synthesis to lower inflammation and oxidative stress (Fig. 4E and Fig. S6). Interestingly, in contrast to the metabolites that were rescued by liver copper supplementation, several different types of carnitines showed increased levels in HFD compared to ND, with further increases in HFD+Cu. Included are the short-chain acylcarnitines, tiglylcarnitine and propionylcarnitine, as well as the long-chain acylcarnitine linoleoylcarnitine, which transports fatty acids of certain length into the mitochondria for fatty acid oxidation (FAO) and lipid breakdown (*51–53*). These data suggest that HFD-induced lipid accumulation stresses cells to enhance FAO, and that copper promotes this process even further (Fig. S7). The SMPDB enrichment analysis further showed alterations in amino acid and energy metabolism as significant features of copper supplementation in HFD compared to HFD alone. Importantly, we observed increases in metabolism of branched-chain amino acids (e.g. valine, leucine, and isoleucine) as well as the malate-aspartate axis, all of which support fatty acid oxidation (Fig. 4F).

Finally, we profiled the liver lipidomes and observed that lipid compositions in liver tissue were altered by copper supplementation in HFD compared to HFD alone (Fig. 4G). The HFD cohorts showed a marked increase in di- and triglycerides, which are primary lipid signatures of steatosis, compared to ND, with liver-targeted copper supplementation effectively lowering these lipids back to normal levels without altering other essential lipids such as phosphatidylcholine (Fig. 4H and I and Table S5). However, this trend was not observed in plasma, iWAT, eWAT, or BAT, indicating that lipid alterations induced by hepatic copper repletion are indeed liver-specific (Fig. S8A-D).

### Copper enhances lipid metabolism by inducing cAMP-dependent lipolysis while suppressing lipid uptake

Proteomics experiments on HFD-induced MASLD mouse livers showed increased expression of PLIN2 and CD36 relative to ND controls, with no changes in other well-characterized lipid droplet markers like PLIN3 and PLIN5. Moreover, hepatic copper supplementation in HFD mice lowered PLIN2 and CD36 expression back to normal levels, while only slightly decreasing PLIN5 (Fig. 5A and B). Interestingly, along these lines we observed rescue of cAMP deficiency in HFD by copper supplementation in mouse MASLD livers (Fig. 5C), suggesting that copper regulates PLIN2 in lipid droplets by a cAMP-mediated signaling mechanism.

**Fig. 5:**
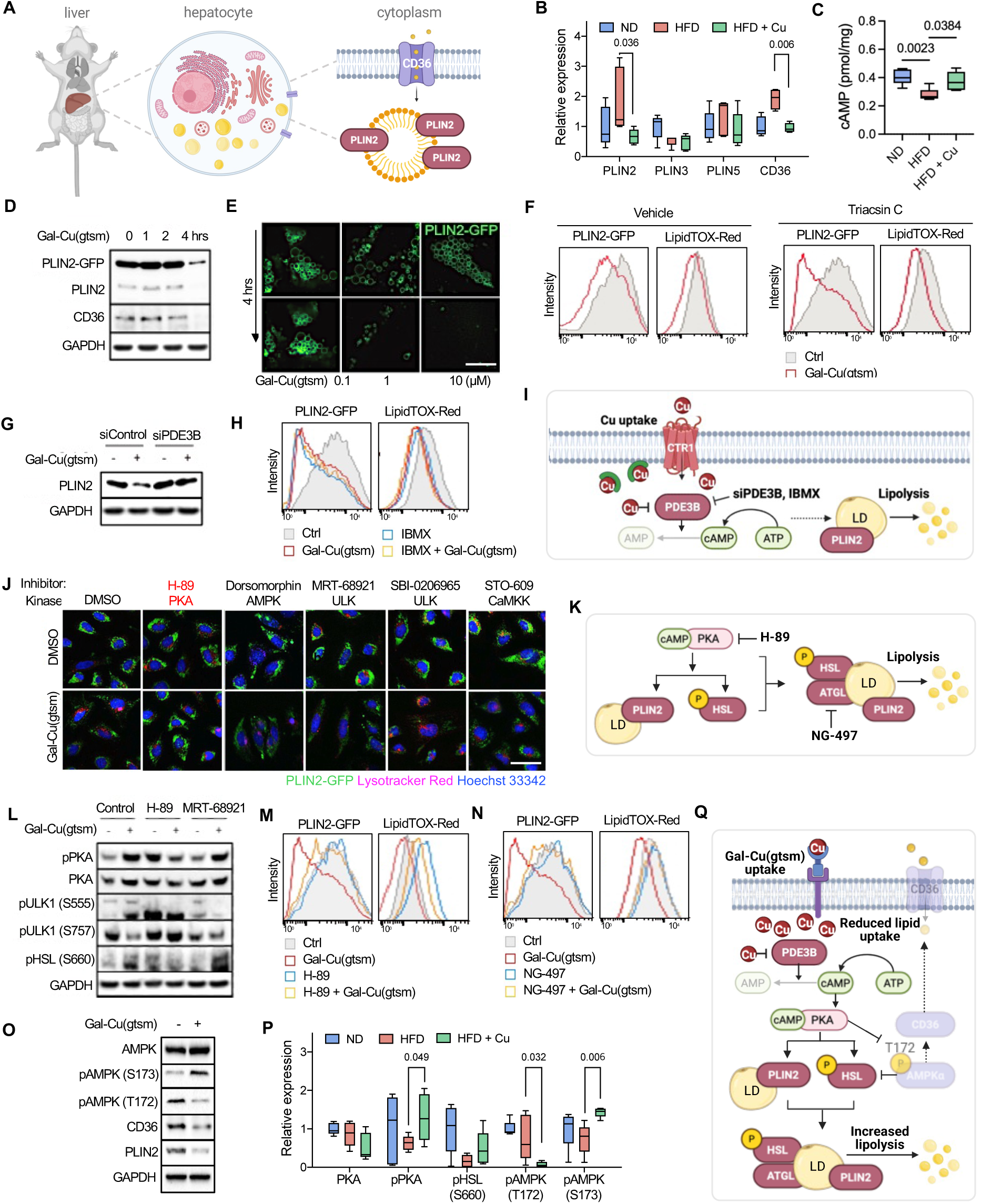
Copper enhances lipid metabolism by inducing cAMP-dependent lipolysis while suppressing lipid uptake. **A.** Overview of PLIN2 and CD36 localization. **B and C.** Mice were fed ND or HFD for 2 weeks followed by Gal-Cu(gtsm) or DPBS vehicle for the following 10 weeks (Fig. 2B, ND, HFD+Cu, n = 6; ND+Cu, HFD, n = 5). **B.** Lipid droplet perilipins (PLINs) and lipid importer CD36 protein expressions in mouse livers. **C.** cAMP measurements in mouse livers. **D.** Immunoblots of Gal-Cu(gtsm) treatment in PLIN2-GFP-Huh7 cells. **E.** Live-cell imaging of PLIN2-GFP-Huh7 cells at 0 and 4 hours with increasing concentrations of Gal-Cu(gtsm). **F.** Flow cytometry histogram of PLIN2-GFP-Huh7 cells pre-treated with vehicle or Triacsin C for lipogenesis inhibition for 24 hours followed by 4 hours of vehicle or Gal-Cu(gtsm) treatment. Cells were measured for GFP for PLIN2 and LipidTOX-Red for non-polar lipids like triglycerides. The scale bar indicates 10 µm. **G.** Immunoblots of PLIN2-GFP-Huh7 cells transfected with scrambled siRNA control or siPDE3B for 24 hours followed by 4 hours of vehicle or Gal-Cu(gtsm) treatment. **H.** Flow cytometry histogram of PLIN2-GFP-Huh7 cells pre-treated with indicated compounds for 24 hours followed by vehicle or Gal-Cu(gtsm) treatment with IBMX phosphodiesterase (PDE) inhibitor for PDE3B. **I.** Schematic of copper supplementation-dependent regulation of lipolysis through PDE3B. Copper, siPDE3B, and IBMX inhibit PDE3B hydrolysis of cAMP, enhancing cAMP-dependent lipolysis **J.** Live-cell imaging of PLIN2-GFP-Huh7 cells treated with inhibitors of kinases potentially affected by copper. Cells were stained with Hoechst 33342 (blue) and Lysotracker (magenta) treated as indicated for 4 hours. The scale bar indicates 50 µm. **K.** Schematic of copper-promoted lipolysis pathway dependent on both PKA and ATGL. **L.** Immunoblots of phosphorylated kinases in PLIN2-GFP-Huh7 cells treated with PKA and ULK inhibitors with copper supplementation for 4 hours. **M and N.** Flow cytometry histogram of PLIN2-GFP-Huh7 cells pre-treated with (M) PKA inhibitor and (N) ATGL inhibitor for 24 hours followed by vehicle or Gal-Cu(gtsm) treatment. **O.** Immunoblots of PLIN2-GFP-Huh7 cells treated with vehicle or 1 µM of Gal-Cu(gtsm) for 24 hours. **P.** Protein expression changes quantified from mouse livers. All error bars represent mean ± SD. Unpaired t-test p-value indicated. **Q.** Proposed mechanism of action of Gal-Cu(gtsm) as a copper supplement to rescue or prevent liver steatosis by copper signaling.

To study biochemical mechanisms in more detail, we moved to examine the effects of copper supplementation with Gal-Cu(gtsm) in a hepatic cell line model of MASLD. We chose Huh7 with stable overexpression of PLIN2-GFP, which exhibits a phenotype with increased lipid droplet accumulation that mimics PLIN2-upregulated MASLD. Upon Gal-Cu(gtsm) treatment, PLIN2-GFP, PLIN2, and CD36 significantly decreased within 4 hours (Fig. 5D). Notably, live-cell imaging experiments demonstrated that not only does copper supplementation with Gal-Cu(gtsm) lower PLIN2 expression, but it also decreases the number and size of PLIN2-containing lipid droplets (Fig. 5E and Fig. S9A). Copper also lowered levels of other lipids as detected by LipidTOX-Red, but to a lesser extent (Fig. 5F). As expected, copper-dependent reduction of PLIN2 was slightly more effective when lipogenesis was blocked with Triacsin C, suggesting that increased lipid synthesis as a compensatory response to Gal-Cu(gtsm) treatment is subtle. To validate this trend, we profiled *de novo* lipogenesis in this Huh7-PLIN2-GFP cell line using metabolic ^13^C-acetate tracing to monitor newly generated lipids labeled with this heavy isotope tag. Consistent with our previous observations in mice liver, copper supplementation in Huh7 noticeably decreased levels of TGs and DGs compared to vehicle control (Fig. S9B). While most are not statistically significant, more TG and DG lipids tended to have reduced labeling from ^13^C-acetate with copper supplementation, indicating slightly reduced *de novo* lipogenesis. Altogether, the results demonstrated that copper mediated lipid regulation was occurring at lipid droplet degradation stage, but not by reducing lipogenesis.

Drawing from metabolomics data that showed HFD-induced cAMP depletion that could be rescued by copper supplementation with Gal-Cu(gtsm) (Fig. 4F), we previously reported that copper can bind to phosphodiesterase 3B (PDE3B) and inhibit its cAMP hydrolase activity by metalloallostery, boosting cAMP-dependent lipolysis in adipocyte models (*20*). We reasoned that copper may play a related signaling role in hepatocytes, where copper-PDE3B metalloallostery represses cAMP hydrolysis, leading to increased cAMP levels and PKA activity to promote downstream PLIN2-containing lipid droplet degradation in MASLD. Indeed, siRNA knockdown of PDE3B blocked copper-triggered downregulation of PLIN2 compared to scrambled control (Fig. 5G). Likewise, treatment with IBMX, a pan-PDE inhibitor, also prevented copper-mediated PLIN2 and lipid droplet degradation (Fig. 5H and I). These data, along with the aforementioned liver metabolomics data showing copper-dependent changes in cAMP levels, establish that copper-PDE3B metalloallostery can regulate hepatocyte lipid droplet metabolism.

Since copper is a nutrient that can modulate several different signaling kinase cascades by direct metal binding to regulate kinase activity, or in the case of PKA, by regulating metabolite substrate levels (*18*, *54–62*), we sought to identify kinase target(s) responsible for copper-mediated PLIN2 degradation by treating hepatic cells with a panel of known inhibitory chemical inhibitors to various kinases with or without Gal-Cu(gtsm) copper supplementation (Fig. 5J and K). Among compounds tested, only H-89, a PKA inhibitor, effectively prevented PLIN2 degradation by copper (Fig. S6C).

With these data in hand establishing a copper-cAMP-PKA signaling axis, we then moved on to study downstream targets in lipid droplet metabolism from both the perspective of lipolysis and lipid uptake. In the context of lipolysis, we found HSL phosphorylation at S660, known to induce lipolysis of lipid droplets upon PKA activation (*63*, *64*), is enhanced by copper treatment, while its copper-dependent phosphorylation was suppressed by co-treatment with a PKA inhibitor H-89 (Fig. 5L). While copper changed phosphorylation of ULK1, another known copper-dependent kinase, at S555 (activating) and S757 (inhibiting) sites, these phosphorylation changes in ULK1, as well as in HSL, remained unchanged when PKA is inhibited by H-89 suggesting that PKA operates at upstream of ULK1. Moreover, ULK1 inhibition by MRT-68921 did not prevent phosphorylation of HSL, demonstrating that copper-activated PKA stimulates HSL lipolysis pathway, independent of ULK1.

As lipolysis of lipid droplets requires PKA-HSL phosphorylation as well as ATGL recruitment, we also assessed copper-dependent lipid regulation upon treatment with PKA and ATGL inhibitors H-89 and NG-497, respectively. We observed that both PKA and ATGL inhibition blocked copper-mediated decreases in lipid levels as measured using PLIN2-GFP by flow cytometry (Fig. 5M and N). Interestingly, because ATGL inhibition alone was not sufficient to increase total lipid levels in contrast to PKA inhibition, the data suggest that a parallel pathway for copper-dependent lipid regulation to reduce total lipid levels without directly affecting PLIN2. Drawing from proteomics data that showed HFD-induced depletion of CD36, a lipid uptake protein, could be rescued by copper supplementation with Gal-Cu(gtsm) (Fig. 4F), we hypothesized that copper could also reduce lipid levels by limiting lipid uptake. Indeed, we observed that Gal-Cu(gtsm) treatment in PLIN2-Huh7 cells also increased the phosphorylation of AMPK, a PKA substrate, preferentially at its S173 site while repressing phosphorylation at its T172 site, the latter of which is known to promote the expression of CD36 (*65*) (Fig. 5O). This result was further validated that phosphorylation of PKA, HSL, and AMPK (S173) was elevated in mouse liver lysates with copper supplementation to support parallel amplification of lipolysis and inhibition of lipid uptake *in vivo* by a copper-cAMP-PKA signaling axis (Fig. 5P). Taken together, we propose a model where liver-targeted copper supplementation with Gal-Cu(gtsm) delivers copper into the cell, where it can inhibit the cAMP hydrolase activity of PDE3B, which subsequently increases cAMP levels and PKA activity to both promote HSL-mediated lipolysis and repress AMPK-mediated CD36 lipid uptake, which work in concert to lower levels of PLIN2-containing lipid droplets (Fig. 5Q).

### Copper rewires mitochondrial energetics toward fatty acid oxidation for lipid degradation

Having shown that copper supplementation promotes lipolysis, we next sought to address how hepatocytes deal with the elevation of free fatty acids that would be produced from increased lipolysis. Since our previous data confirmed that copper supplementation improves liver health (Fig. 2C, F, G and Fig. 3B-I) and restores liver lipid metabolism without largely altering lipid profiles in plasma, adipose tissue, or skeletal muscle (Fig. S2D-F), we hypothesize that enhanced mitochondrial fatty acid oxidation in liver could provide a pathway for the metabolism of free fatty acids (Fig. 6A). Along these lines, analysis of metabolomic and proteomic data revealed a landscape of copper-dependent changes that are FAO-associated. Long-chain acyl-carnitines for mitochondrial fatty acid transport were significantly increased by Gal-Cu(gtsm) copper supplementation, as well as 3-aminoisobutyrate, associated with fat browning and increased FAO (*66–68*) (Fig. 4E and 6B). Moreover, proteomics data showed that Gal-Cu(gtsm) copper supplementation upregulated many protein targets in fatty acid oxidation, including the key FAO enzyme CPT2 (Fig. 6C). Unlike PLIN2 or CD36, which were elevated in HFD and decreased back by copper treatment, the expression levels of FAO proteins were not affected by diet alone, suggesting that this effect was not solely a result of improved liver health, but that copper exhibits a beneficial effect of enhancing FAO selectively under conditions of excess lipids (Table S1 and S3).

**Fig. 6:**
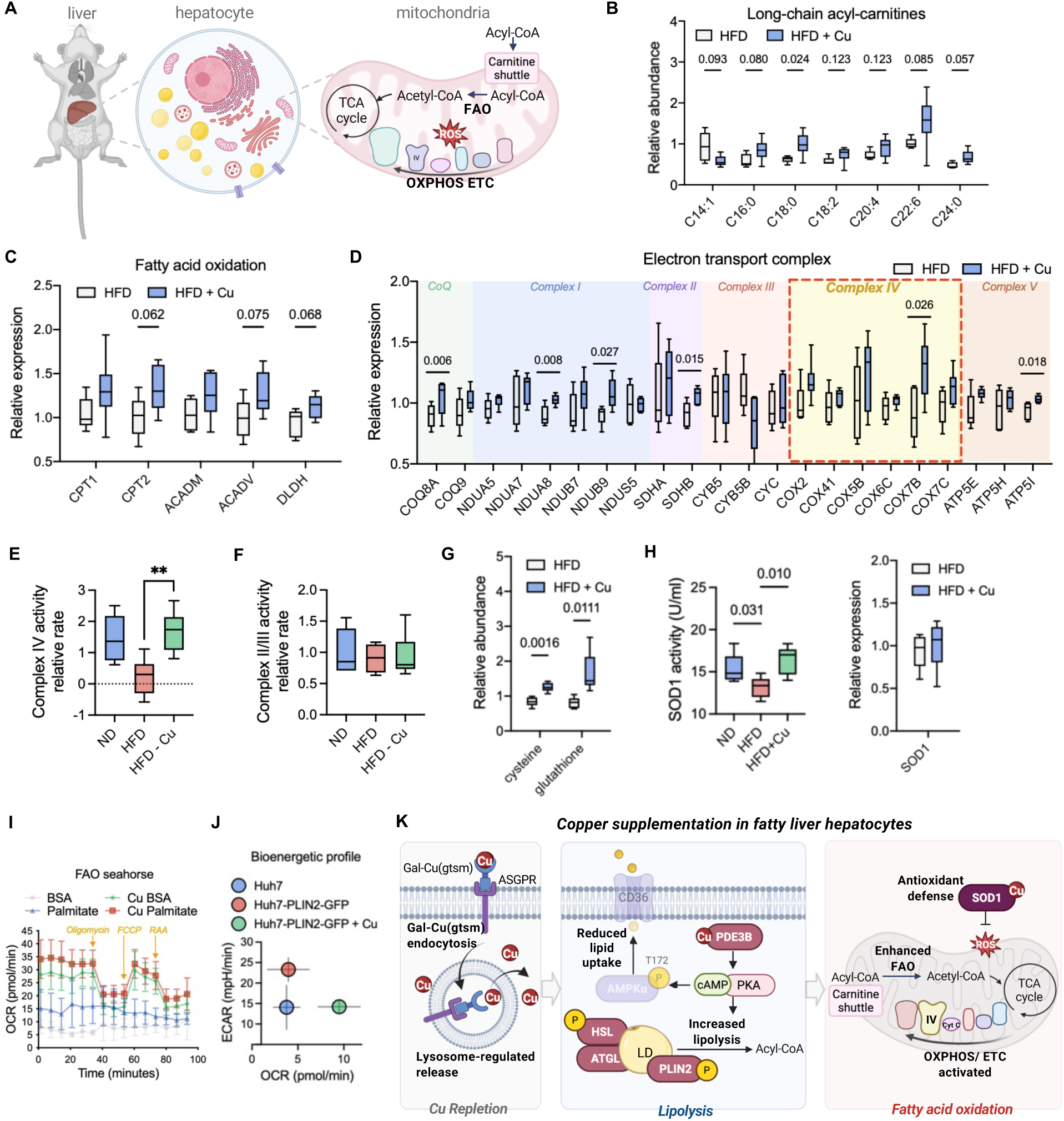
Copper rewires mitochondrial energetics toward fatty acid oxidation for lipid degradation. **A.** Overview of mitochondria metabolism in liver that was studied. **B-H.** Mice were fed with ND or HFD for 2 weeks followed by Gal-Cu(gtsm) or DPBS for the following 10 weeks (Fig. 2B, ND, HFD+Cu, n = 6; ND+Cu, HFD, n = 5). **B.** Relative abundance of fatty acid oxidation (FAO)-associated metabolites measured in mouse livers. **C.** Relative protein expression of FAO-associated proteins. **D.** Relative protein expression of mitochondria electron transport complex organized by their Complex. Complex IV (red box) is copper-dependent. **E and F.** Relative ETC complex activity measured in mitochondria suspension isolated from mouse livers. **G.** Relative abundance of antioxidant-associated metabolites. **H.** SOD1 activity measured from mouse livers. **I.** BSA-palmitate Seahorse assay of OCR with copper supplementation. Cells were treated with or without Gal-Cu(gtsm) for 24 hours followed by palmitate or BSA for 15 minutes before measuring OCR and ECAR. **J.** Seahorse assay of extracellular acidification rate (ECAR) for glycolysis and oxygen consumption rate (OCR) for mitochondrial respiration in Huh7 and PLIN2-GFP-Huh7 cells treated with or without Gal-Cu(gtsm) for 24 hours. **K.** Combined proposed mechanism of Gal-Cu(gtsm) liver-targeted supplementation and the roles of copper in inducing lipolysis and enhancing fatty acid oxidation in fatty liver disease. All error bars represent mean ± SD. All error bars represent mean ± SD. Unpaired t-test; ns, not significant, ∗p < 0.05, ∗∗p < 0.01, ∗∗∗p < 0.001, ∗∗∗∗p < 0.0001 or exact numbers indicated.

Several targets in the electron transport chain (ETC) across complex I, II, IV and V subunits, NDUA8, NDUB9, SDHB, COX7B, and ATP5I, were significantly upregulated in Gal-Cu(gtsm)-treated HFD livers compared to control HFD livers and many of the other protein expression trended upwards (Fig. 6D), pointing towards increased mitochondrial oxidative capacity. Strikingly, copper supplementation restored complex IV activity while complex II/III remained unaffected (Fig. 6E and F), presumably due to the requirement for copper as a nutrient cofactor for both complex IV assembly and activity (*69*, *70*). As further evidence for elevated oxidative capacity, we observed increased antioxidant response. Metabolites that are responsive to reactive oxygen species (ROS) responses, such as cysteine and glutathione, increased in copper supplemented livers (Fig. 6G and Fig. S7F) (*71*). Moreover, while expression was not changed, the deficiency in activity of the main intracellular antioxidant SOD1, a copper-dependent enzyme, was rescued by Gal-Cu(gtsm) supplementation (Fig. 6H), strengthening redox capacity to support increased mitochondrial metabolism.

The collective data suggest that hepatic copper supplementation by Gal-Cu(gtsm) not only restores ETC complex IV and SOD1 antioxidant defense activity as a nutrient cofactor but also rewires mitochondrial metabolism to effectively protect cells against excess lipid accumulation. For functional evidence whether copper treatment induces metabolic rewiring toward fatty acid oxidation, we performed a BSA-palmitate Seahorse assay in Huh7 hepatoma-derived cells (Fig. 6I). In vehicle-treated cells, the addition of palmitate elicited only a modest increase oxygen consumption rate (OCR) relative to the BSA-only control, indicating that Huh7 cells exhibit low basal mitochondrial respiration, which is consistent with a more glycolytic, cancer-derived metabolism. In contrast, copper supplementation with Gal-Cu(gtsm) markedly increased OCR relative to vehicle-treated controls, with palmitate slightly elevating OCR above BSA-only. Copper elicited robust responses to mitochondrial perturbations with oligomycin, FCCP, and rotenone-antimycin A (RAA), whereas vehicle-treated cells exhibited minimal OCR. Consistent with this increased FAO capacity and with the previously shown protein expression changes in mouse livers, Gal-Cu(gtsm) treatment in Huh7 cells upregulated CPT2, and to a lesser extent CPT1 and cytochrome c oxidase subunit 1 (COX1) (Fig. S10A).

In contrast, glycolytic flux, as assessed by ECAR, was not significantly perturbed by copper (Fig. S10B). Indeed, glycolytic and TCA intermediates (Fig. S10C) and citrate synthase activity as a proxy for TCA activity (Fig. S10D) were largely unaffected by HFD feeding or Gal-Cu(gtsm) treatment in mouse livers. In addition, we further examined how bioenergetic profiles in hepatic cell models change with copper supplementation. PLIN2-GFP overexpression shifted basal metabolism of Huh7 cells toward glycolysis as shown by increasing extracellular acidification (ECAR) (Fig. 6J). Since PLIN2 stabilized lipid droplets resulting in decreased fatty acid oxidation, it is reasonable that glycolysis would increase upon PLIN2 overexpression. The addition of Gal-Cu(gtsm) decreased ECAR back to the basal levels and increased OCR, indicating reduced glycolysis back to levels of cells with endogenous levels of PLIN2 expression and increased mitochondrial respiration. Here, copper supplementation has the potential to support the adaptation towards elevated mitochondrial respiration and restoration of the metabolic imbalance in MASLD. Taken together, these results support the model that copper supplementation reprograms metabolism specifically toward mitochondrial respiration and enhanced fatty acid oxidation capacity, providing a mechanistic explanation for its ability to reduce lipid accumulation in copper-supplemented livers (Fig. 6K).

## Discussion

Here, we identified dysregulated copper homeostasis as a nutrient vulnerability in a large combined MASLD and MASH patient RNA-seq dataset and in CDAHFD-induced MASH mouse models, consistent with prior observations of copper deficiency in high-fat diet (HFD)-fed mice (*24*). To interrogate the functional roles of copper in steatotic liver disease, we applied our Targeted Ionophore-based Metal Supplementation (TIMS) strategy (*35*), using Gal-Cu(gtsm) to enable liver-directed copper delivery as a potential therapeutic approach for addressing copper-related pathologies. Using this platform, we showed that Gal-Cu(gtsm) improves steatosis in HFD-fed mice, conferring both preventive and restorative effects, while simultaneously lowering blood glucose levels without compromising lean mass or skeletal muscle integrity.

To characterize the molecular and physiological consequences of hepatic copper supplementation in MASLD models, we integrated transcriptomic, proteomic, metabolomic, and lipidomic datasets. Through assessment of significant changes across the different layers of omics datasets, we observed that HFD-induced lipid dysregulation is marked by upregulation of key lipid droplet (PLIN2) and lipid uptake (CD36) proteins, which was reversed by Gal-Cu(gtsm)-mediated copper supplementation. Using a PLIN2-GFP–expressing Huh7 cell model, we further established that copper inhibits phosphodiesterase 3B (PDE3B) cAMP hydrolase activity, leading to increased levels of cAMP and subsequent activation of protein kinase A (PKA). This signaling cascade promotes phosphorylation of HSL at S660, selectively targeting PLIN2-marked lipid droplets for lipolysis. In parallel, copper-mediated PKA activation preferentially induces phosphorylation of AMP-activated protein kinase at S173, with a lesser effect at T172, resulting in suppression of CD36 expression and reduced lipid uptake.

Beyond lipid droplet turnover, we traced how hepatocytes process the excess free fatty acids generated by copper-induced lipolysis. Copper supplementation with Gal-Cu(gtsm) reprograms cellular metabolism toward mitochondrial respiration through upregulation of electron transport chain (ETC) complex IV activity. This metabolic shift enables enhanced fatty acid oxidation (FAO) while maintaining redox balance, as excess reactive oxygen species (ROS) generated through and oxidative phosphorylation (OXPHOS) are effectively buffered by activation of copper-dependent superoxide dismutase 1 (SOD1). Together, these findings reveal that copper acts as a central regulator that coordinates lipid mobilization, lipid uptake suppression, and mitochondrial oxidative metabolism to resolve diet-induced lipid accumulation in hepatocytes.

Collectively, our results demonstrate both prophylactic and therapeutic effects of hepatic copper supplementation in MASLD and establish a mechanistic framework by which copper reprograms lipid and energy metabolism to counter steatosis. Still, this study has some limitations. While we observed selective hepatic copper deficiency preceding the onset of steatosis, the cause of this copper depletion is not clear. Finding the cause of copper deficiency may provide further insight for disease prevention strategies. In addition, our study reveals robust metabolic benefits of hepatic copper repletion, it does not directly address fibrotic and inflammatory processes that characterize more advanced stages of disease. From the striking copper homeostasis changes we observed in human MASH samples and CDAHFD diet-induced mouse model of liver fibrosis, there is evidence that copper also plays roles in livers that have progressed to inflammation and fibrosis. As studies continue to reveal roles of copper in cellular regulation, investigations of the relationship between copper and fibrosis progression, as well as development into hepatocellular carcinoma will further enable translational applications.

## Methods

### Synthesis of copper supplements

The liver-targeted copper supplement Gal-Cu(gtsm) and untargeted Cu(gtsm) control supplement were synthesized by previously reported methods (*35*, *72*).

### Meta-analysis of MASLD and MASH patient data

GSE48452, GSE61260, GSE83452, GSE167523, GSE33814, GSE89632, GSE136261, GSE126848, GSE130970, and PRJNA512027 dataset was combined and analyzed for top differential genes and gene ontology enrichment analysis in R following Piras’s paper.(*26*) Briefly, raw data were preprocessed to get normalized data adjusted for surrogate variables. Then we used the *GeneMeta* R package to standardize data using z-score transformation. The FDR for each gene was generated using the *ZscoreFDR* function with 50,000 permutations, and FDR < 0.05 was considered to be statistically significant for differential gene expression. These DEGs were put into *enrichGO* function for gene ontology analysis and p-values were adjusted using FDR method.

### Cell culture

All cells were maintained as a monolayer in Dulbecco’s Modified Eagle Medium + GlutaMAX (DMEM, Gibco) supplemented with 10% (v/v) fetal bovine serum (FBS, Seradigm) at 37 °C in a 5% CO_2_ atmosphere.

### Cytotoxicity measurements

Huh7 cells were grown to 80% confluency in 100 µL complete media in a 96-well plate at 37 °C, 5% CO_2_. Cells were then treated with Gal-Cu(gtsm) or Cu(gtsm) in DMSO for a range of concentrations (0, 1, 10 µM) in multiplicate and incubated at 37 °C, 5% CO_2_ for 4 hours. Following incubation, the media was then aspirated and replaced with 100 µL fresh phenol-free media containing 10 µM DCFDA (Abcam; ab113851) and 5 µM MitoSOX (Abcam; ab219943), or 10 µM CalceinAM and 5 µg/ml 7-AAD (Abcam; ab270789). After 1 hour of incubation, fluorescence intensity was read by plate reader at 535 nm (DCFDA) and 595 nm (MitoSOX) or at 517 nm (CalceinAM) and 647 (7-AAD).

### CDAHFD-induced MASH model mice

Mice were fed a choline-deficient, L-amino acid-defined, high-fat diet (CDAHFD) that consists of 60% kcal% fat and 0.1% methionine by weight to induce a MASH mouse model.10 Mice were killed, and livers were extracted, flash frozen, and stored at –80 °C until analysis. Serum was collected by coagulating blood samples for 1 hour at RT, centrifuging at 1500 × g for 15 min at 4 °C, flash freezing, and storing at –80 °C. CDAHFD-fed mice (n = 5) were compared to normal diet-fed mice (n = 5).

### Metal measurements in livers by ICP-MS

About 30 mg of frozen liver were cut on dry ice for each sample and resuspended in 1 mL HNO3/100 mg tissue. Samples were diluted, and ^63^Cu, ^65^Cu, ^56^Fe, and ^57^Fe were measured using a Thermo Fisher iCAP-Qc ICP-MS in KED mode and reported in ppm or normalized by liver weight.

### Copper supplementation in mice

All animal studies were approved by and performed according to the guidelines of the Animal Care and Use Committee of the University of California, Berkeley. Mice were group housed on a 12:12 hour light-dark cycle at 22 °C with free access to food and water.

For the prophylactic studies that demonstrate copper prevents lipid accumulation, C57BL male mice were fed with high-fat diet (60% kcal% fat, Research Diets D12492) or normal diet (10% kcal% fat, sucrose matched with D12492, Research Diets D12450J) for 2 weeks. Copper in these diets were normalized based on caloric density, containing 6.5–7 mg copper per 4057 kcal. Then mice were supplemented with copper by intraperitoneal injection of 7.26 mg Gal-Cu(gtsm)/kg mouse—equivalent to 0.75 mg Cu/kg mouse—twice per week for 10 weeks while continuing their respective diets. Mice were sacrificed after the combined 12 weeks, and tissue samples were collected and stored in –80 °C for the 4 groups (ND, HFD+Cu, n = 6; ND+Cu, HFD, n = 5).

For the therapeutic studies that demonstrate copper reduces lipids after accumulation, C57BL male mice were fed with high-fat diet (60% kcal% fat, Research Diets D12492) or normal diet (10% kcal% fat, sucrose matched with D12492, Research Diets D12450J) for 12 weeks. Copper in these diets were normalized based on caloric density, containing 6.5–7 mg copper per 4057 kcal. Then mice were supplemented with copper by intraperitoneal injection of 7.26 mg Gal-Cu(gtsm)/kg mouse—equivalent to 0.75 mg Cu/kg mouse—twice per week for 8 weeks while getting normal diet. Mice were sacrificed after the combined 20 weeks, and tissue samples were collected and stored in –80 °C for the 4 groups (n = 6).

### Oil Red O staining of tissue slides

For visualization of lipid droplets in ND, ND+Cu, HFD, and HFD+Cu liver samples, liver sections were stained with Oil Red O (Abcam; ab150678). The frozen tissue section slides were incubated in propylene glycol for two minutes, in Oil Red O solution for 6 minutes, differentiated in 86% propylene glycol for 1 minute, and then rinsed twice in water. Then the slides were incubated in hematoxylin for 2 minutes, rinsed three times in water, mounted with aqueous mounting medium and coverslip, and imaged using microscope.

### Liver health measurements

ALT and AST enzyme activity were measured from mouse plasma using commercial kits (Invitrogen; EEA001 and EEA003) following manufacturer’s protocols. Briefly, the plasma samples were centrifuged at 10000 g for 10 minutes and 5 µL of the supernatant was reacted with 20 µL of substrate solution and 20 µL of chromogenic agent to be incubated at 37 °C for 20 minutes. Then 200 µL of alkali reagent was added, mixed with microplate reader for 10 seconds, and let stand for 15 minutes at room temperature before the OD value being measured at 510 nm. For standard curve, 0, 2, 4, 6, or 8 mmol/L sodium pyruvate was reacted with 20, 18, 16, 14 or 12 µL of substrate solution instead of plasma. From the OD, blank OD was subtracted, and the activity (IU/L) was calculated using the standard curve. Measurements of CHOL, HDL, LDL, ALT activity, and AST activity were performed at Merck & Co., Inc., Rahway, NJ, USA by using 50 µL of serum were diluted 2X with PBS and run on their Clinical Analyzer.

### RT-qPCR of MASLD-associated genes

RNA was handled with appropriate precautions with RNase-free materials and RNaseZap Decontamination Solution (Invitrogen). 5–10 mg of frozen liver tissue was used per sample to extract RNA with the PureLink RNA Mini Kit (Invitrogen) following the kit’s protocol for animal tissue. Briefly, samples were homogenized with a handheld pestle motor and passing the lysate through a 25-gauge needle. Lysates were taken through the protocol and eluted with 30 µL RNase-free water. Sample concentrations were measured by NanoDrop and then digested with DNase I. A 1% agarose gel was run to evaluate RNA quality. Reverse transcription was performed with 1.4 µg RNA per sample using the High-Capacity cDNA Reverse Transcription Kit (Applied Biosystems), using a ratio of 400 ng RNA in a 20-µL reaction. All samples were pooled for a no reverse transcriptase control. qPCR was performed on the cDNA using TaqMan Fast Advanced Master Mix on the Bio-Rad CFX96 Real-Time PCR System (Applied Biosystems) protocols for 96-well plates. Made-to-order TaqMan Array (FAM), format 16, 96-well plates were designed with the 16 assays for mouse listed below. Each cDNA sample was run in triple replicate per gene target. No reverse transcriptase and no template controls were also run. The 2–ΔΔCT method was used to calculate normalized expression change.

MASLD-associated genes that were analyzed by RT-qPCR with their respective TaqMan Gene Expression Assay IDs. * Indicate reference genes.

| Gene | Gene name | Assay ID |
| --- | --- | --- |
| * *18S* | Eukaryotic 18S rRNA | Hs99999901_s1 |
| * *Hprt* | Hypoxanthine guanine phosphoribosyl transferase | Mm00446968_m1 |
| * *Ywhaz* | Tyrosine 3-monooxygenase/tryptophan 5-monooxygenase activation protein zeta | Mm03950126_s1 |
| * *Rplp0* | Ribosomal protein lateral stalk subunit P0 | Mm00725448_s1 |
| *Srebf1* | Sterol regulatory element binding transcription factor 1 | Mm00550338_m1 |
| *Nr1h3* | Nuclear receptor subfamily 1; group H; member 3 | Mm00443451_m1 |
| *Fasn* | Fatty acid synthase | Mm00662319_m1 |
| *Acaca* | Acetyl-Coenzyme A carboxylase alpha | Mm01304257_m1 |
| *Scd1* | Stearoyl-Coenzyme A desaturase 1 | Mm00772290_m1 |
| *Dgat2* | Diacylglycerol O-acyltransferase 2 | Mm00499530_m1 |
| *Elovl5* | Elongation of long chain fatty acids, family member 5 | Mm00506717_m1 |
| *Nfe2l2* | Nuclear factor, erythroid derived 2, like 2 | Mm00477784_m1 |
| *Pnpla3* | Patatin-like phospholipase domain containing 3 | Mm00504420_m1 |
| *Srebf2* | Sterol regulatory element binding factor 2 | Mm01306292_m1 |
| *Scap* | SREBF chaperone | Mm01250176_m1 |
| *Ccl2* | Chemokine (C-C motif) ligand 2 | Mm00441242_m1 |

### Proteomics of mouse liver tissue

TMT10plex Mass Tag Labeling Kit reagents and protocol were used to prepare peptides for protein quantification. Frozen liver tissue was lysed with the kit’s Lysis Buffer following the method discussed above. 100 µg of lysate was reduced, alkylated, precipitated by methanol/chloroform, and digested with Trypsin/Lys-C Mix. Mass Spec Grade (Promega) overnight following the manufacturer’s protocol. Peptide concentration was measured using Pierce Quantitative Colorimetric Peptide Assay (Thermo Scientific), and 20 µg peptide was labeled with 0.2 mg TMT label in a total volume of 46 µL. Three sets of TMT10plex were prepared with 8 samples and 2 pooled samples each. Pooled samples were from all 24 samples, 6 samples from the 4 groups. Samples were quenched with 2 µL 5% hydroxylamine. The three sets of 10 labeled samples were combined into 1.5 mL tubes and vacuum dried. The three tubes of TMT10plex-labeled peptides were submitted to University of California, Davis, Proteomics Core Facility for proteomic profiling with tandem mass spec analysis.

### Metabolomics

At least 100 mg frozen liver for each mice were analyzed using the Metabolon HD4 metabolomics platform, operated by Metabolon. The metabolite samples were prepared to remove proteins and separate into fractions to analyze using different MS methods: two different reverse phases (RP)/UPLC-MS/MS with positive ion mode ESI, RP/UPLC-MS/MS with negative ion mode ESI, and HILIC/UPLC-MS/MS with negative ion mode ESI. Methods utilize a Waters ACQUITY UPLC and Thermo Scientific Q Exactive MS. Metabolite data was extracted using Metabolon’s methods to obtain area-under-the-curve for each feature.

### Lipidomics

Lipid samples were prepared by tissue extraction using 1:1 DCM:methanol overnight at 4 °C followed by a modified Bligh-Dyer extraction using methanol/water/DCM with deuterated internal standards. The extracts were then dried under nitrogen and reconstituted in 250 µL of 10 mM ammonium acetate in 50:50 DCM:methanol. MS analysis was performed on a Shimazdu LC with nano PEEK tubing and Sciex SelexIon-5500 QTRAP in both positive and negative ion mode electrospray ionization (ESI). Individual lipid species were quantified in nmol/g tissue by taking the peak area ratio of the species with their respective deuterated internal standard, then multiplying the concentration of internal standard added to the sample. Water extract blanks were subtracted from sample quantification as background, lipid class, lipid classes, lipid species, fatty acid concentrations were calculated.

### De novo lipogenesis labeling

For ^13^C-acetate labeling, Huh7-PLIN2-GFP cells were cultured in 12-well plates with or without 1 µM of Gal-Cu(gtsm) in presence of 5 mM acetate or ^13^C-acetate in high glucose DMEM with 10 % FBS for 24 hours. The media was collected into another plate placed on dry ice. Next, 250 µL pre-cooled IPA was added on each well with cells or with pure media without any cells or treatment for blank. The metabolites were extracted into 1.5 mL tubes and spun 21,000 x g for 20 minutes at 4 °C. The supernatant was removed and placed into new tubes and spinning was repeated once more. The final supernatant was collected in new tubes and 100 µL of sample was loaded into pre-chilled mass spec vials with inserts. The same procedure was performed using the media to analyze lipid secretion. Then 5 µL of each sample was injected on LC-MS and the results were analyzed using El-Maven (*73*), natural abundance was corrected, PCA plot drawn for determining any outliers and shifts between groups, and log2FC and log10(p-value) was calculated in R for plotting.

### Statistical tests and analysis of omics datasets

Data was normalized by dividing each value by the median of the feature (i.e., lipid species, metabolite, or protein) across all samples, so that the median for each feature is equal to 1. Features with any missing values were filled with the minimum value across all samples. The processed values were then saved as a CSV file and analyzed by using the web interface of MetaboAnalyst 5.0 (https://www.metaboanalyst.ca) or in R studio. The datasets were analyzed by PCA, and two sample outliers were identified and removed based on the clustering. The data was then analyzed by ANOVA and Tukey HSD to identify changes with significant changes. For visualization, data was first filtered for significant changes between HFD and ND. These filtered features were then analyzed by volcano plot for significant changes between HFD+Cu and HFD and that the direction of the change was opposite from the change between HFD and ND.

### Immunofluorescence imaging

Huh7-PLIN2-GFP cells were grown to 90% confluency in 300 µL complete media in 8-well ibdis chamber at 37 °C, 5% CO_2_. For live-cell imaging, cells were treated with Gal-Cu(gtsm) in DMSO for a range of concentrations (0.1, 1, 10 µM) in duplicates. Right after adding treatments, cells were imaged at eight distinct fields per condition at 20x magnification using Zeiss LSM confocal microscope and Zen software. After 4 hours of incubation, cells were imaged at the saved positions. The PLIN2-GFP intensity was measured and quantified using FIJI software. For representative images, cells were imaged at 63x magnification.

### Kinase inhibition assays

For kinase inhibition assay, cells were treated with 5 µM of Gal-Cu(gtsm) or DMSO with or without each kinase inhibitor as below for 3 hours. After incubation, cells were washed with PBS and labeled with 50 nM Lysotracker deep red (Invitrogen; L12492) and 5 µg/mL Hoechst 33342 (Invitrogen; #62249) in PBS for 30 minutes. Cells were washed two times in PBS and imaged in fresh PBS at eight distinct fields per condition at 20x magnification using Zeiss LSM confocal microscope and Zen software.

| Item/Compounds | CAS | Final conc. |
| --- | --- | --- |
| STO-609 (acetate), CaMKK1/2 inhibitor | 1173022-21-3 | 1 μg/mL |
| H-89 (hydrochloride), PKA inhibitor | 130964-39-5 | 5 μM |
| MRT68921, ULK1/2 inhibitor | 1190379-70-4 | 1 μM |
| SBI-0206965, ULK1/2 inhibitor (also AMPK) | 1884220-36-3 | 10 μM |
| Dorsomorphin, non-selective AMPK inhibitor | 866405-64-3 | 5 μM |

For immunoblotting, Huh7-PLIN2-GFP cells were treated with H-89, MRT68921 or DMSO vehicle for 3 hours with or without 5 µM Gal-Cu(gtsm), trypsinized, and pelleted to be stored at −80 °C until before lysis.

### siRNA knock-down

Huh7-PLIN2-GFP cells were grown to 70% confluency in 3 mL complete media in a 6-well plate at 37 °C, 5% CO_2_. Transfection was then performed as per Lipofectamine RNAiMax protocol. Briefly, 150 pmol of desired siRNA, siControl (Thermo; 4390843) or siPDE3B (Thermo; AM16708), was introduced to 7.5 µL lipid. The lipid-siRNA complex was incubated for 30 minutes at 23 °C in 500 µL Opti-MEM media (Gibco). Then, 500 µL complex was added to 2.5 mL complete media. Next, 10 mL of this solution was added to each 10 cm plate containing a final siRNA concentration of 50 nM. Cells were incubated for 24 hours at 37 °C, 5 % CO_2_. The following day, cells were harvested by trypsinization. Cells were washed 1X with 1 mL of ice-cold PBS, pelleted at 300 xg, and flash-frozen in LN_2_ for future analysis.

### Flow cytometry

Huh7-PLIN2-GFP cells were grown to 70% confluency in 3 mL complete media in a 6-well plate at 37 °C, 5% CO_2_. Cells were treated with 5 µM Triacsin C, 1 µM of IBMX, or 1 µM of NG-497 for 24 hours for inhibiting lipogenesis, PDE3B, or ATGL, if desired, prior to Gal-Cu(gtsm) 5 µM or DMSO, and co-treated with 5 µM of H-89 for PKA inhibition for 6 hours. Then the cells were washed with PBS, trypsinized, and resuspended in 5 µM LipiTox-Red (Invitrogen; H34476) for 30 minutes. The cells were spun down for 3 minutes at 300 x g, resuspended in fresh PBS, and analyzed using BD FACS analyzer.

### Immunoblotting of proteins

Approximately 5 mg of frozen liver were homogenized using a handheld pellet pestle with 10 µL lysis buffer/1 mg liver tissue using RIPA Lysis and Extraction Buffer (25 mM Tris-HCl pH 7.6, 150 mM NaCl, 1% NP-40, 1% sodium deoxycholate, 0.1% SDS, Thermo Scientific) with cOmplete Protease Inhibitor Cocktail with EDTA (Roche). For cell samples, cell pellets were vortexed every 10 minutes in same lysis buffer for 30 minutes on ice. Homogenized lysates were incubated on ice for 30 minutes and centrifuged at 12,000 × g for 20 min, 4 °C. Lysates were pipetted while avoiding the lipid layer at the top and transferred to a new tube. Protein concentration was quantified using Pierce BCA Protein Assay Kit (Thermo Fisher). 20 - 40 µg of protein lysate boiled with 4X Laemmli’s buffer (Bio-Rad; 1610747) + 10% BME diluted at 95 °C for 8 minutes. Approximately 30 µg of protein sample was loaded per well and separated on precast 4–20% Novex Tris-Gly SDS-PAGE gels (Invitrogen) run at 160 V for 70 minutes. Proteins were then electro-transferred to a PVDF membrane (25 V/2.5 A for 30 minutes). Blots were blocked in 5% milk in TBS-Tween for 1 hour at RT then incubated in primary antibodies at various dilutions in 5% BSA in TBS-Tween overnight on a shaker at 4 °C. Blots were then incubated with their respective secondary antibodies at 1/3000 dilution in 5% BSA in TBS-Tween for 1 hour at RT. ECL western blotting substrates (Sigma; 34580) for 5 minutes and imaged with ChemiDoc MP were used to image the blots. Blots were normalized to GAPDH loading control using the Bio-Rad Image Lab software.

Primary and secondary antibodies used for immunoblotting. * Indicate antibodies that have been discontinued.

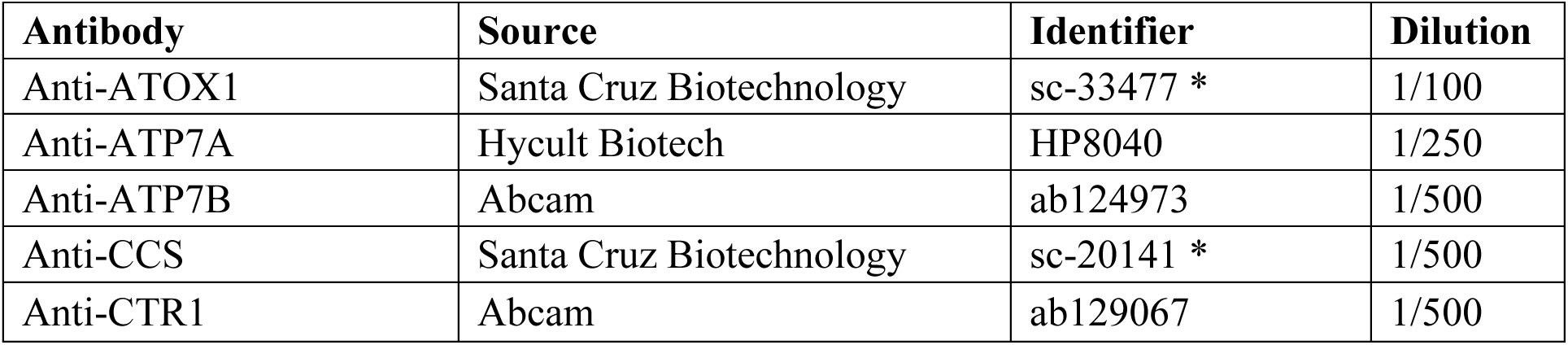
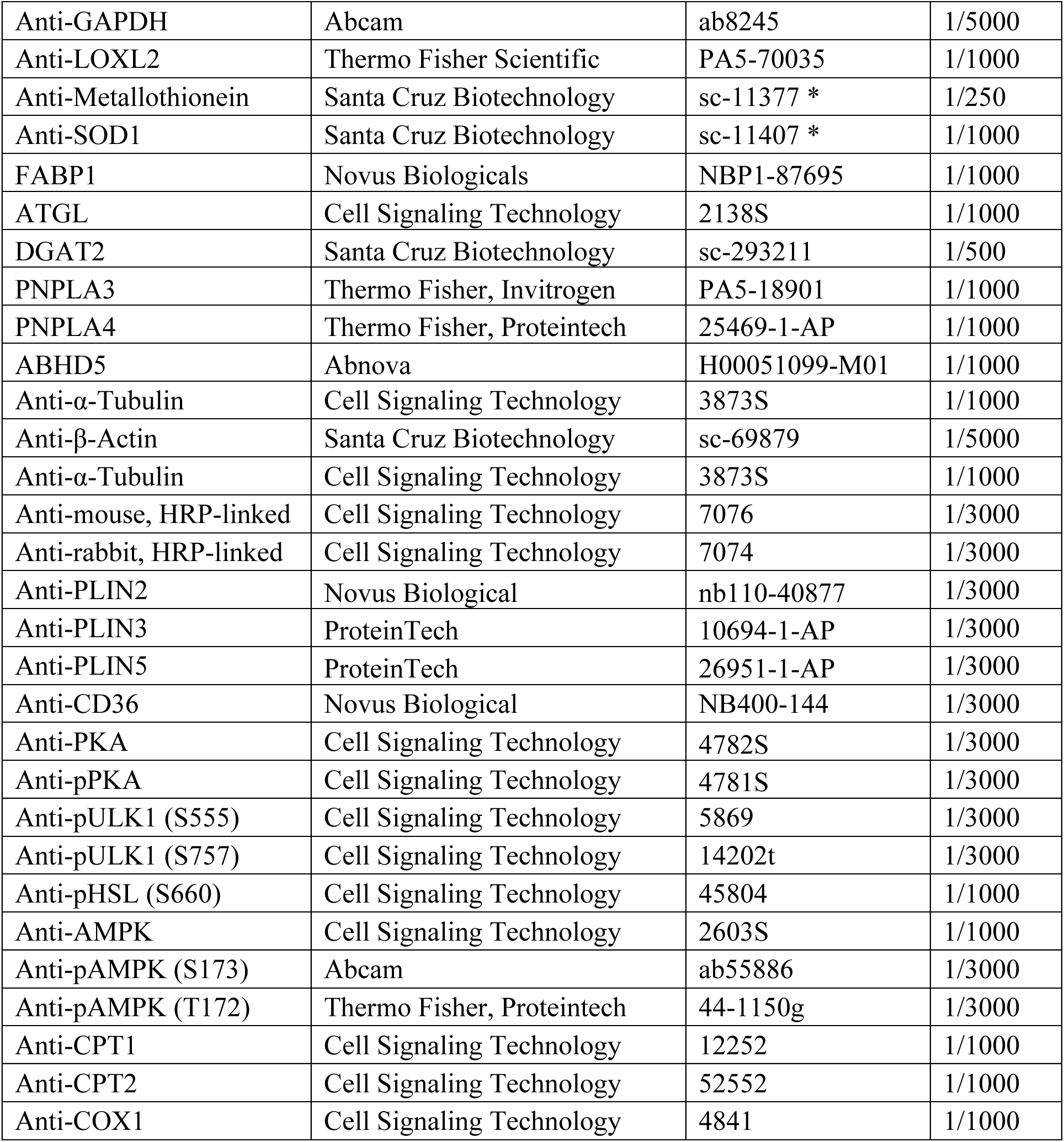

| Antibody | Source | Identifier | Dilution |
| --- | --- | --- | --- |
| Anti-ATOX1 | Santa Cruz Biotechnology | sc-33477 * | 1/100 |
| Anti-ATP7A | Hycult Biotech | HP8040 | 1/250 |
| Anti-ATP7B | Abcam | ab124973 | 1/500 |
| Anti-CCS | Santa Cruz Biotechnology | sc-20141 * | 1/500 |
| Anti-CTR1 | Abcam | ab129067 | 1/500 |
| Anti-GAPDH | Abcam | ab8245 | 1/5000 |
| Anti-LOXL2 | Thermo Fisher Scientific | PA5-70035 | 1/1000 |
| Anti-Metallothionein | Santa Cruz Biotechnology | sc-11377 * | 1/250 |
| Anti-SOD1 | Santa Cruz Biotechnology | sc-11407 * | 1/1000 |
| FABP1 | Novus Biologicals | NBP1-87695 | 1/1000 |
| ATGL | Cell Signaling Technology | 2138S | 1/1000 |
| DGAT2 | Santa Cruz Biotechnology | sc-293211 | 1/500 |
| PNPLA3 | Thermo Fisher, Invitrogen | PA5-18901 | 1/1000 |
| PNPLA4 | Thermo Fisher, Proteintech | 25469-1-AP | 1/1000 |
| ABHD5 | Abnova | H00051099-M01 | 1/1000 |
| Anti-α-Tubulin | Cell Signaling Technology | 3873S | 1/1000 |
| Anti-β-Actin | Santa Cruz Biotechnology | sc-69879 | 1/5000 |
| Anti-α-Tubulin | Cell Signaling Technology | 3873S | 1/1000 |
| Anti-mouse, HRP-linked | Cell Signaling Technology | 7076 | 1/3000 |
| Anti-rabbit, HRP-linked | Cell Signaling Technology | 7074 | 1/3000 |
| Anti-PLIN2 | Novus Biological | nb110-40877 | 1/3000 |
| Anti-PLIN3 | ProteinTech | 10694-1-AP | 1/3000 |
| Anti-PLIN5 | ProteinTech | 26951-1-AP | 1/3000 |
| Anti-CD36 | Novus Biological | NB400-144 | 1/3000 |
| Anti-PKA | Cell Signaling Technology | 4782S | 1/3000 |
| Anti-pPKA | Cell Signaling Technology | 4781S | 1/3000 |
| Anti-pULK1 (S555) | Cell Signaling Technology | 5869 | 1/3000 |
| Anti-pULK1 (S757) | Cell Signaling Technology | 14202t | 1/3000 |
| Anti-pHSL (S660) | Cell Signaling Technology | 45804 | 1/1000 |
| Anti-AMPK | Cell Signaling Technology | 2603S | 1/1000 |
| Anti-pAMPK (S173) | Abcam | ab55886 | 1/3000 |
| Anti-pAMPK (T172) | Thermo Fisher, Proteintech | 44-1150g | 1/3000 |
| Anti-CPT1 | Cell Signaling Technology | 12252 | 1/1000 |
| Anti-CPT2 | Cell Signaling Technology | 52552 | 1/1000 |
| Anti-COX1 | Cell Signaling Technology | 4841 | 1/1000 |

### Mitochondrial purification

For mitochondria isolation, approximately 100 mg of frozen livers were washed with ice-cold PBS twice, added with 800 µL of BSA/Reagent A solution (Invitrogen; MAN0011535) in gentleMACS C Tubes (Miltenyi Biotec; #130-093-237) and homogenized in gentleMACS Octo Dissociator (Miltenyi Biotec; #130-095-937) by running “m_mito_tissue_01” program. Then 800 µL of Isolation Reagent C (Invitrogen; MAN0011535) was added, inverted several times to mix, and centrifuged at 700 x g for 10 minutes at 4 °C. The supernatant was transferred into a new tube, centrifuged again at 6000 x g for 15 minutes at 4 °C, and the supernatant cytosolic fraction was saved for SOD1 activity test. The mitochondrial pellet was washed with 400 µL of Wash Buffer followed by centrifugation at 12000 x g for 5 minutes and resuspended in fresh 200 µL of Wash Buffer. Both cytosolic and mitochondrial extracts were stored at −80 °C until used.

### Mitochondrial enzymatic activity tests

Isolated mitochondrial was diluted 1:200 in assay media provided by complex II/III (Cayman; #700950), complex IV (Cayman; #700990) and citrate synthase (Cayman; #701040) activity assay kits. In each kit, 6 ND, 6 HFD, and 6 HFD+Cu liver mitochondrial fractions were analyzed following manufacturer’s protocol.

Briefly, complex II/III activity assay measured complex III-dependent reduction of cytochrome c coupled to complex II by adding 50 µL of assay buffer A (958 µL assay buffer + 20 µL bovine heart mitochondria reagent + 2 µL of 1 mM rotenone and 20 µL of 100 mM KCN) and 20 µL of samples to 30 µL of reagent B (reduced cytochrome c and succinate assay reagent diluted in assay buffer for 1:10 and 1:100, respectively) in a 96-well assay plate. The plate was immediately placed in plate reader to measure absorbance at 550 nm at 30 second internals for 15 minutes at 25 °C.

Complex IV activity was assayed by measuring the oxidation rate of reduced cytochrome c was by adding 50 µL of assay buffer A (995 µL assay buffer + 5 µL bovine heart mitochondria reagent) and 20 µL of samples to 30 µL of reagent B (1:10 diluted reduced cytochrome c assay reagent in assay buffer) in a 96-well assay plate. The plate was immediately placed in plate reader to measure absorbance at 550 nm at 30 second internals for 15 minutes at 25 °C.

Citrate synthase activity was measured by adding 50 µL of assay buffer A (acetyl-CoA and developer reagent both diluted in assay buffer by 1:50) and 20 µL of samples to 30 µL of reagent B (oxaloacetate reagent diluted in assay buffer by 1:25) in a 96-well assay plate. The plate was immediately placed in plate reader to measure absorbance at 412 nm at 30 second internals for 20 minutes at 25 °C. For comparison, the maximum reaction rate (Vmax) calculated for each sample and compared between the groups.

### SOD1 activity assay

The cytosolic fraction from mitochondrial isolation was used for SOD1 activity measurements using commercial kit (Abcam; DIASODC). Tissue lysates were diluted by 1:4 in assay buffer. Briefly, 10 µL of standards (92, 46, 23, 11.5, 5.8, 2.9, 1.4 or 0 U/mL) or diluted samples were added with 50 µL of substrate solution (250 µL of substrate concentrate in 2.25 mL xanthine oxidase buffer) and 25 µL of xanthine oxidase solution (50 µL of xanthine oxidase in 1.2 mL xanthine oxidase buffer) in a 96-well assay plate. The plate was incubated for 20 minutes at room temperature and was placed in plate reader to measure absorbance at 450 nm. For the blank control, plate was read at 450 nm before adding xanthine oxidase. The absorbance values from blank control were subtracted from corresponding absorbance measured for each sample, and the activity was calculated using the standard curve.

### Seahorse assays

Seahorse assays were performed using Agilent kits and the XFe analyzer instrument. Huh7-PLIN2-GFP and Huh7 cells were plated in XFe 96-well plate (Agilent; 103792-100) in DMEM with 10 % FBS. When cells reached 85% confluency, cells were treated with 1 µL of Gal-Cu(gtsm) or DMSO for 24 hours. The day before assay, a sensor cartridge was hydrated in their Calibrant solution (Agilent; 100840-000) in a non-CO_2_ incubator. On the day of assay, assay medium was prepared by supplementing Seahorse XF DMEM (Agilent; 103575-100) with 1 mM pyruvate (Agilent; 103578-100), 2 mM glutamine (Agilent; 103579-100), and 10 mM glucose (Agilent; 103577-100). Then 20 µL oligomycin, 22 µL FCCP, and 25 µL RAA (Agilent; 103015-100) were placed into port B, C and D of sensor cartridge to make final 1.5 µM, 0.5 µM and 0.5 µM, respectively, when added to the well. The media in the cell plate was changed to assay media and cells were stabilized in non-CO_2_ incubator for 40 minutes while sensor cartridge was calibrated in XFe analyzer. For the palmitate assay, 30 µL of BSA or BSA-Palmitate solution (Agilent; 102720-100) was added to each well right before assay started. The ECAR and OCR values were measured by MitoStress program and analyzed using Wave software.

## Author contributions

J.K., V.N.P., A.S., and C.J.C. conceived of this study. T.A.S., W.C., and Y.A. synthesized the chemical compounds used in this study. J.K analyzed patient metadata. T.A.S., I.L., and D.S. performed mouse copper supplementation experiments, measured blood glucose and EchoMRI, and harvested tissues. T.A.S. and T.X. performed ICP-MS. A.T.P. performed cell imaging assays. C.N. and S.T. provided CDAHFD liver samples, clinical liver health panel measurements, resources, and advice and expertise on MASLD. J.D.R. provided resources for mass spectrometry and metabolomics/lipidomics. J.K. and V.N.P. performed the multi-omic analyses. J.K. performed and analyzed cellular assays, mitochondrial isolation and enzymatic assays based on discussions with J.O., I.L., and X.X. The manuscript was written by J.K., V.N.P. and C.J.C. and all authors contributed to the editing of this manuscript.

## Acknowledgements

We thank Dr. A. Killilea and her staff from the UC Berkeley Cell Culture Facility for technical support with cell culture. We thank Dr. Yang Liu, Dr. Xueyuan Jiang, and Anu Sebin from Merck & Co., Inc., Rahway, NJ, USA for technical support on liver and serum measurements. We thank Christopher Novotny and Saswata Talukdar for helpful discussions on MASLD mouse models and feeding experiments. We thank Dr. Kimberly Jackson and Dr. Priya Ramamoorthy from Metabolon for support with metabolomics and lipidomics. We thank Dr. Mike Lange and Dr. Melissa Roberts from UC Berkeley for helpful discussions on lipid metabolism and Prof. Marie Heffern from UC Davis for helpful advice on diet studies. We thank Prof. Donita Brady, Dr. Jaclyn Welles and Dr. Paul M. Titchenell for helpful discussions on feeding experiments and fibrosis mouse models. Schematics were created with BioRender.

## Supplementary Materials

Supplemental Materials and Methods

Table S1 (Source data and image quantification)

Table S2-5 (Multi-omics raw data)

Table S6 (Multi-omics enrichment analysis)

---

**Fig. S1:**
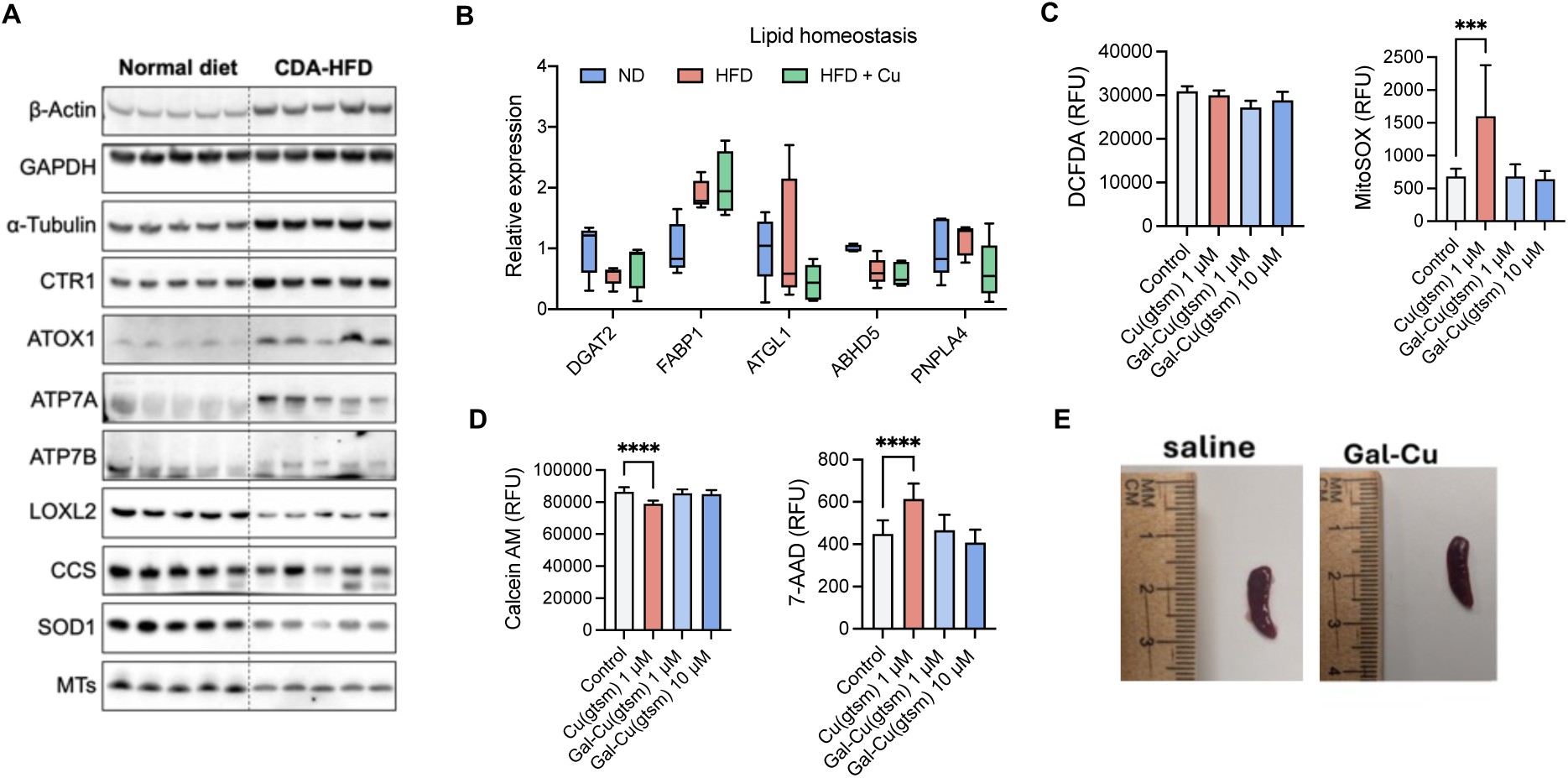
Copper is depleted in fatty livers while copper supplementation protects liver against fat accumulation. **A.** Immunoblots of copper homeostasis markers in ND and CDAHFD livers. **B and E.** Mice were fed with ND or HFD for 2 weeks followed by Gal-Cu(gtsm) or DPBS for the following 10 weeks (ND, HFD+Cu, n = 6; ND+Cu, HFD, n = 5). **B.** Relative expression of lipid homeostasis in livers. **C and D.** Comparison of toxicity of Cu(gtsm) with liver-targeted Gal-Cu(gtsm) by (D) DCFDA cellular ROS assay and mitochondrial superoxide (MitoSOX) and (E) live (Calcien AM) and dead (7-AAD) cells measured after indicated treatment for 4 hours in Huh7 cells. **E.** Representative image of spleen to check for copper treatment toxicity from mice treated with HFD for 2 weeks followed by Gal-Cu(gtsm) or DPBS vehicle for the following 10 weeks. All error bars represent mean ± SD. Unpaired t-test; ns, not significant, ∗p < 0.05, ∗∗p < 0.01, ∗∗∗p < 0.001, ∗∗∗∗p < 0.0001 or exact numbers indicated.

**Fig. S2:**
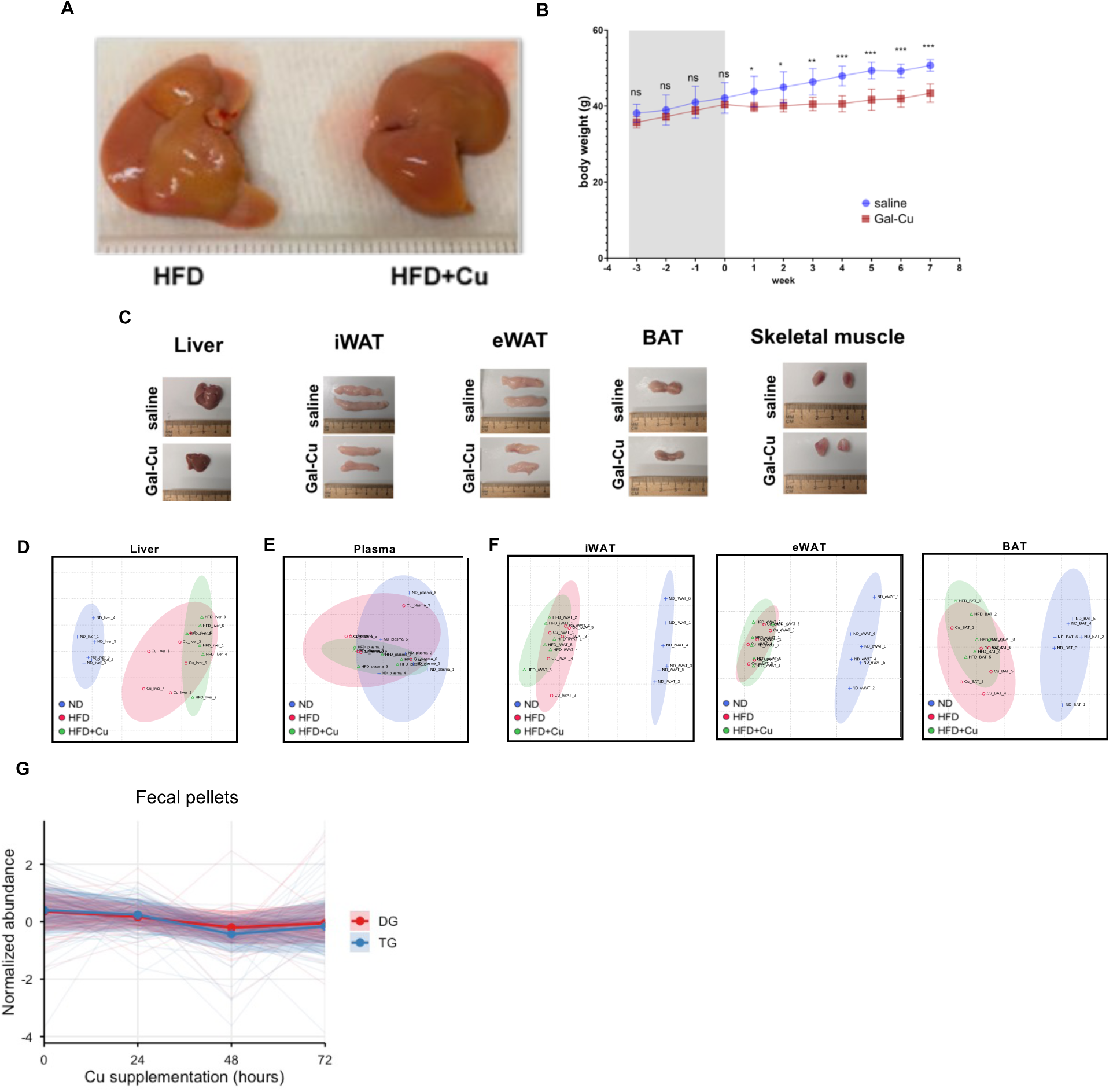
Copper supplementation in steatotic livers alter physiological phenotypes in mice. **A.** Representative image of livers from mice fed with HFD for 2 weeks followed by Gal-Cu(gtsm) or DPBS for the following 10 weeks. **B-G.** Mice were fed normal diet or HFD for 12 weeks then normal diet for 8 weeks treatment with Gal-Cu(gtsm) or DPBS (n = 6). **B.** Mouse body weight measured before and during 8 weeks during copper supplementation for HFD-fed mice. Week 0 is the start of Gal-Cu(gtsm) treatment **C.** Representative images of tissues at the end point of therapeutic study of HFD-diet fed mice. **D-F.** PCA of di- and triglyceride lipid profiles in (D) liver, (E) plasma, and (F) adipose tissues. **G.** Normalized di- and triglyceride abundance in fecal pellets with 0, 24, and 48, and 72 hours of copper supplementation. Each light-colored line represents a lipid species. The solid line and shaded region represent mean ± SD of the DGs and TGs. All error bars represent mean ± SD. Anova; ns, not significant, ∗p < 0.05, ∗∗p < 0.01, ∗∗∗p < 0.001, ∗∗∗∗p < 0.0001 or exact numbers indicated.

**Fig. S3:**
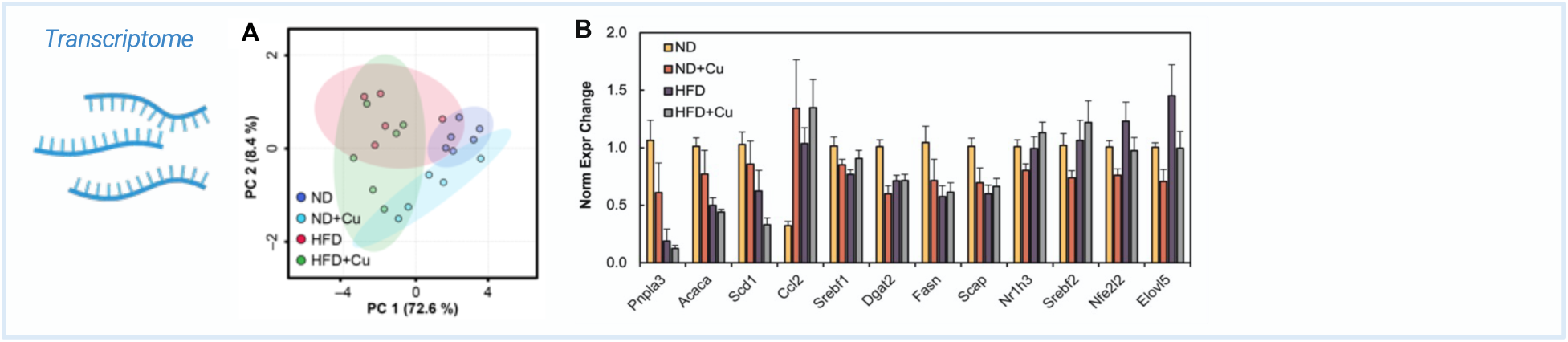
RNA transcripts of HFD and copper supplementation. **A-B.** Analysis of liver from mice that were fed with ND or HFD for 2 weeks followed by treatment with Gal-Cu(gtsm) or DPBS vehicle for the following 10 weeks (Fig. 2B, ND, HFD+Cu, n = 6; ND+Cu, HFD, n = 5 per group). **A.** PCA of mouse liver RT-qPCR transcripts. **B.** RT-qPCR of 12 MASLD-associated transcripts.

**Fig. S4:**
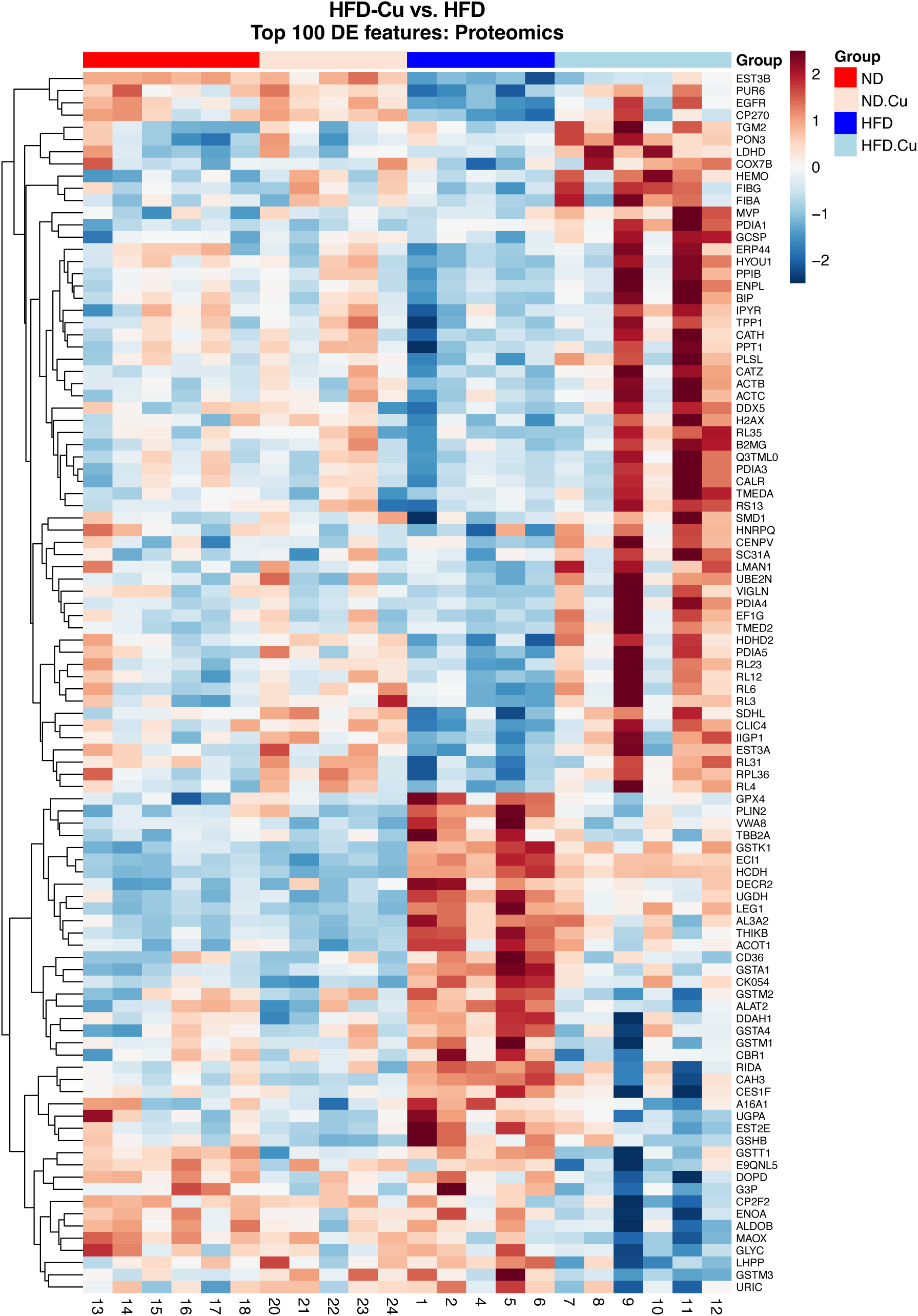
Heatmap of proteomics top 100 DEG comparing HFD+Cu vs. HFD. The liver proteome from mice fed with ND or HFD for 2 weeks followed by Gal-Cu(gtsm) or DPBS for the following 10 weeks (ND, HFD+Cu, n = 6; ND+Cu, HFD, n = 5) were analyzed. Heatmap shows top 100 proteins ordered by HFD+Cu vs. HFD. The color bar represents z-scores.

**Fig. S5:**
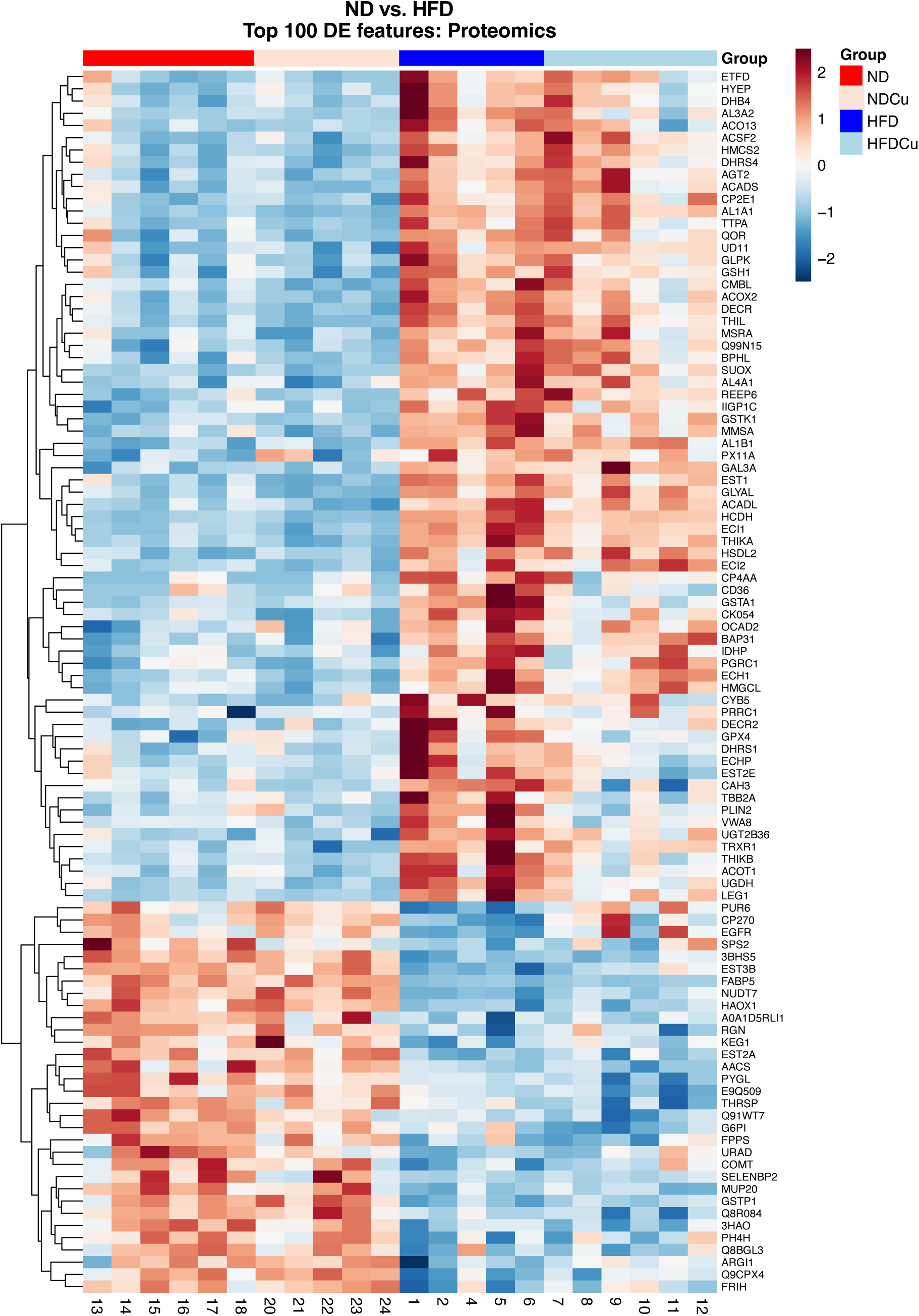
Heatmap of proteomics top 100 DEG comparing HFD vs. ND. The liver proteome from mice fed with ND or HFD for 2 weeks followed by Gal-Cu(gtsm) or DPBS for the following 10 weeks (ND, HFD+Cu, n = 6; ND+Cu, HFD, n = 5) were analyzed. Heatmap shows top 100 proteins ordered by ND vs. HFD. The color bar represents z-scores.

**Fig. S6:**
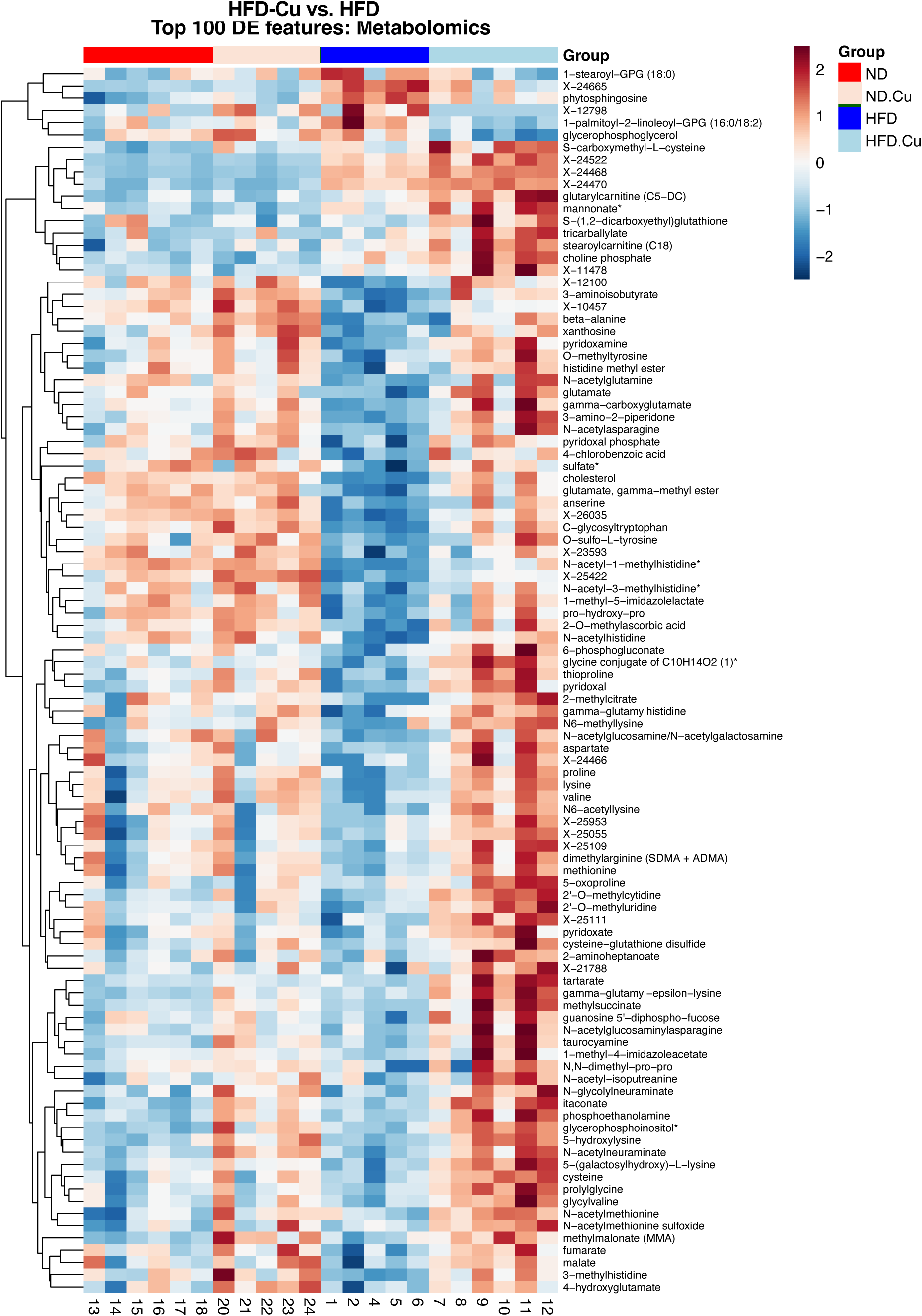
Heatmap of metabolomics top 100 DEG comparing HFD+Cu vs. HFD. The liver metabolomes from mice fed with ND or HFD for 2 weeks followed by Gal-Cu(gtsm) or DPBS for the following 10 weeks (ND, HFD+Cu, n = 6; ND+Cu, HFD, n = 5) were analyzed. Heatmap shows top 100 metabolites ordered by HFD+Cu vs. HFD. The color bar represents z-scores.

**Fig. S7:**
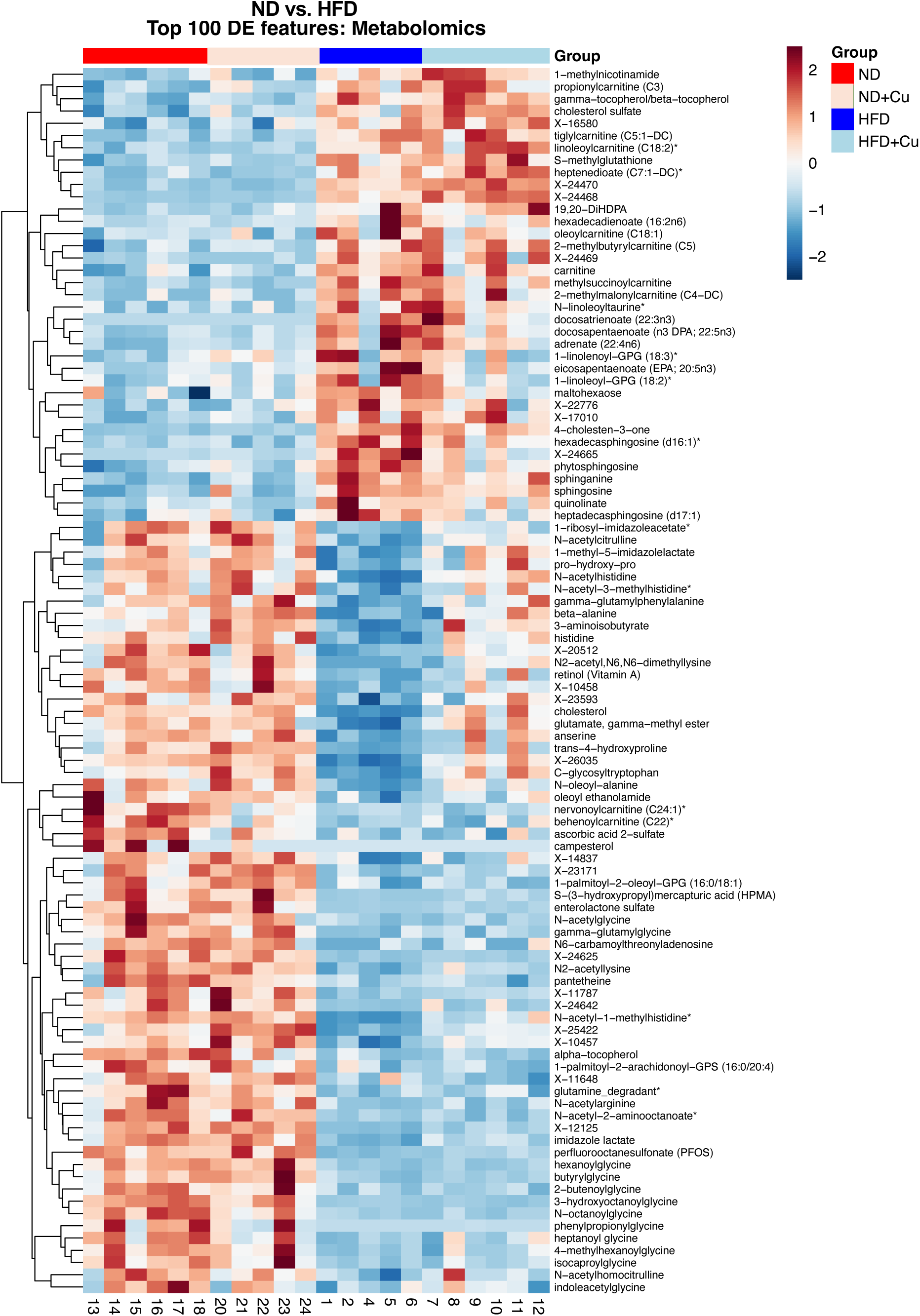
Heatmap of metabolomics top 100 DEG comparing HFD vs. ND. The liver metabolomes from mice fed with ND or HFD for 2 weeks followed by Gal-Cu(gtsm) or DPBS for the following 10 weeks (ND, HFD+Cu, n = 6; ND+Cu, HFD, n = 5) were analyzed. Heatmap shows top 100 metabolites ordered by ND vs. HFD. The color bar represents z-scores.

**Fig. S8:**
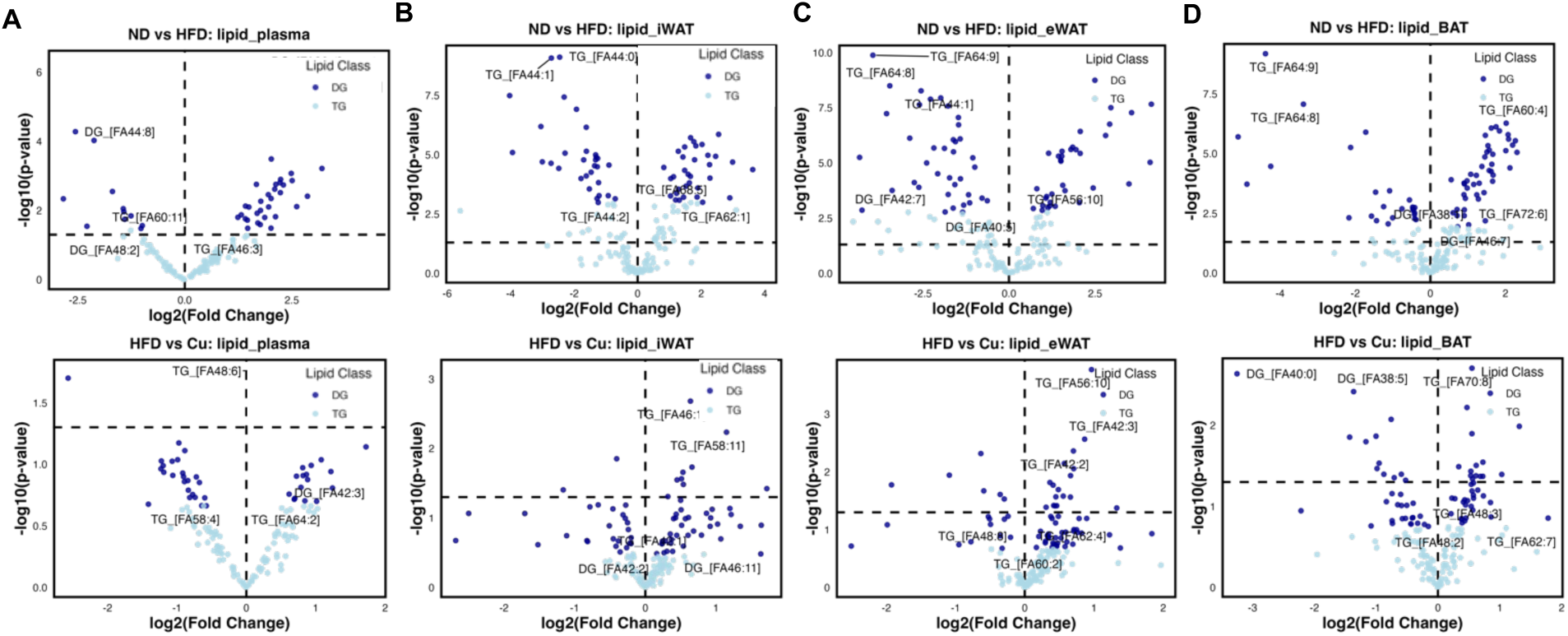
Lipidomics of liver-neighboring adipose tissues and plasma. **A-D.** The tissues from mice fed with ND or HFD for 12 weeks followed by Gal-Cu(gtsm) or DPBS for the following 8 weeks with ND were collected for analyzing lipidomics, and the volcano plots drawn with TG and DG classes. **A.** Volcano plots of ND vs. HFD and HFD vs. Cu plasma lipidomics. **B.** Volcano plots of ND vs. HFD and HFD vs. Cu iWAT lipidomics. **C.** Volcano plots of ND vs. HFD and HFD vs. Cu eWAT lipidomics. **D.** Volcano plots of ND vs. HFD and HFD vs. Cu BAT lipidomics.

**Fig. S9:**
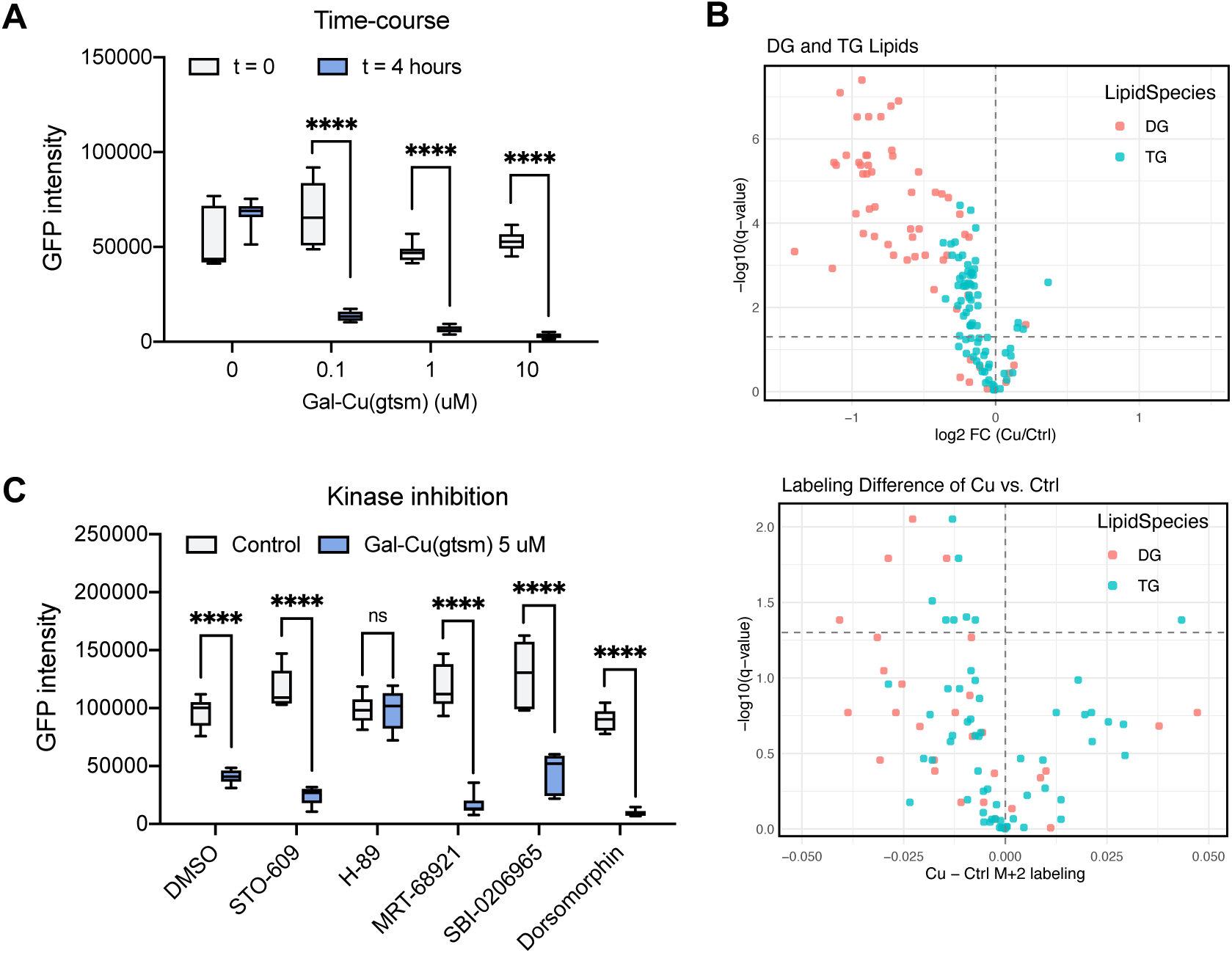
Copper effects metabolome and lipidome in hepatocytes. **A.** Live-cell imaging of PLIN2-GFP-Huh7 cells traced for 4 hours after indicated treatment. **B.** De novo lipogenesis of PLIN2-GFP-Huh7 cells. Cells were incubated with unlabeled acetate (top) or heavy-acetate (bottom) for 24 hours for labeling followed by Gal-Cu(gtsm) treatment for 4 hours and analyzed. **C.** Live-cell imaging of PLIN2-GFP-Huh7 cells stained with Hoechst 33342 (blue) and lysotracker (red) after treatment for 4 hours as indicated.

**Fig. S10:**
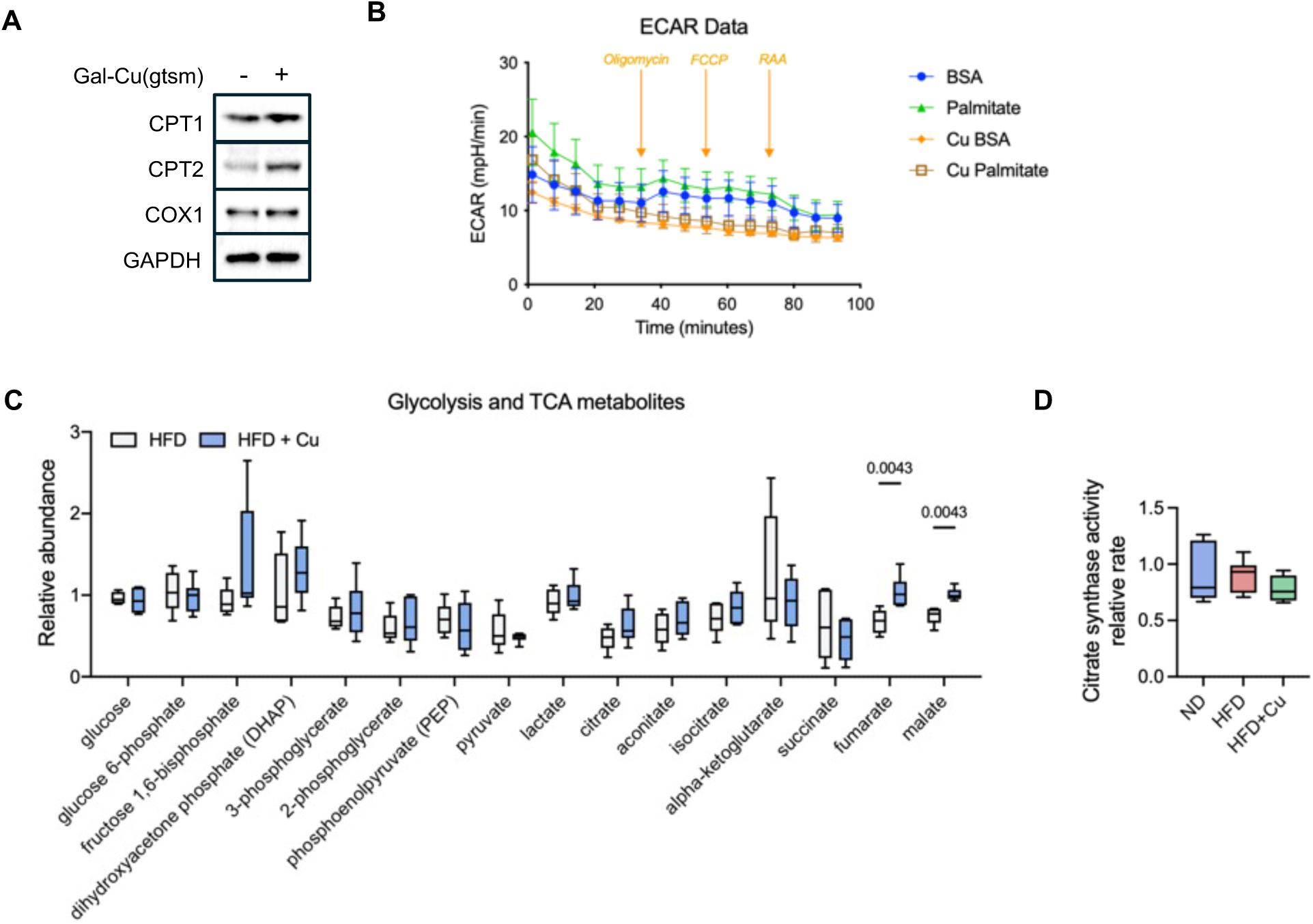
Copper effects metabolome and lipidome in hepatocytes. **A.** Immunoblotting of Huh7 cells treated with 1 mM of Gal-Cu(gtsm) or DMSO for 24 hours. **B.** ECAR measurement from seahorse assays on Huh7 cells after 4 hours of pretreatment of vehicle or Gal-Cu(gtsm). All error bars represent mean ± SD. **C.** The relative metabolite abundance and protein expression levels in liver from mice fed with ND or HFD for 2 weeks followed by Gal-Cu(gtsm) or DPBS for the following 10 weeks (ND, HFD+Cu, n = 6; ND+Cu, HFD, n = 5) measured. **D.** Relative enzyme activity measured in mitochondria suspension isolated from mouse livers from mice fed with ND or HFD for 2 weeks followed by Gal-Cu(gtsm) or DPBS for the following 10 weeks (ND, HFD+Cu, n = 6; ND+Cu, HFD, n = 5).

