## Supplementary Figures for "A Liver-Targeted Copper Supplement Reduces Metabolic Dysfunction-Associated Liver Steatosis by Increasing Lipolysis and Fatty Acid Oxidation"

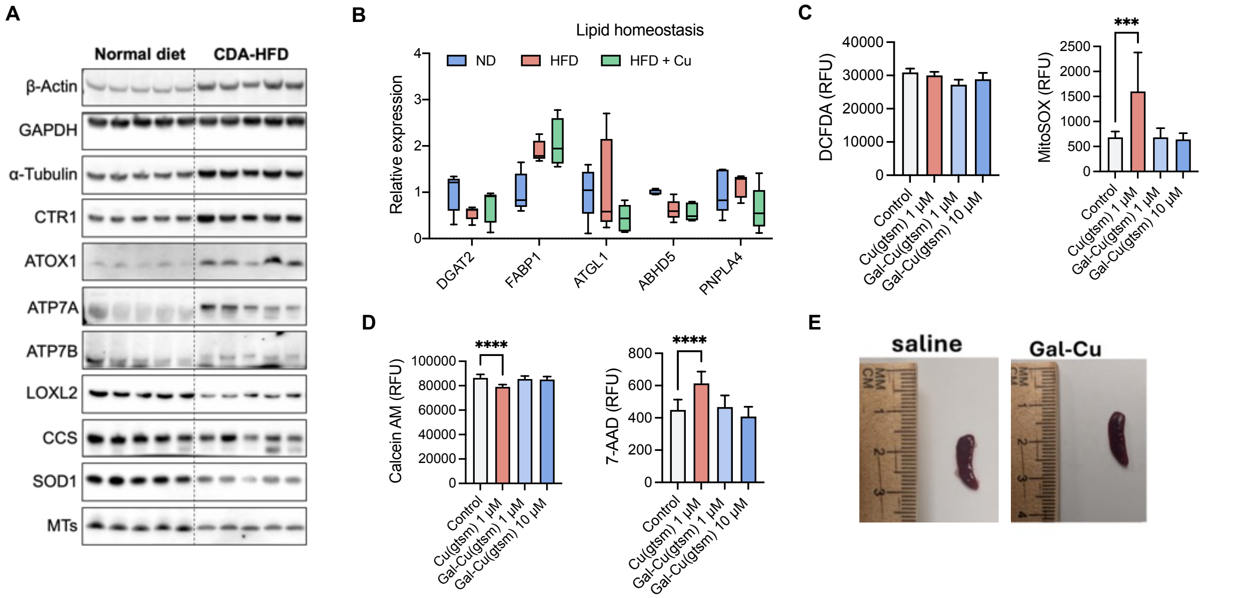


**Fig. S1: Copper is depleted in fatty livers while copper supplementation protects liver against fat accumulation.**

**A.** Immunoblots of copper homeostasis markers in ND and CDAHFD livers. **B and E.** Mice were fed with ND or HFD for 2 weeks followed by Gal-Cu(gtsm) or DPBS for the following 10 weeks (ND, HFD+Cu, n = 6; ND+Cu, HFD, n = 5). **B.** Relative expression of lipid homeostasis in livers. **C and D.** Comparison of toxicity of Cu(gtsm) with liver-targeted Gal-Cu(gtsm) by (D) DCFDA cellular ROS assay and mitochondrial superoxide (MitoSOX) and (E) live (Calcien AM) and dead (7-AAD) cells measured after indicated treatment for 4 hours in Huh7 cells. **E.** Representative image of spleen to check for copper treatment toxicity from mice treated with HFD for 2 weeks followed by Gal-Cu(gtsm) or DPBS vehicle for the following 10 weeks. All error bars represent mean ± SD. Unpaired t-test; ns, not significant, $*p < 0.05$, $**p < 0.01$, $***p < 0.001$, $****p < 0.0001$ or exact numbers indicated.

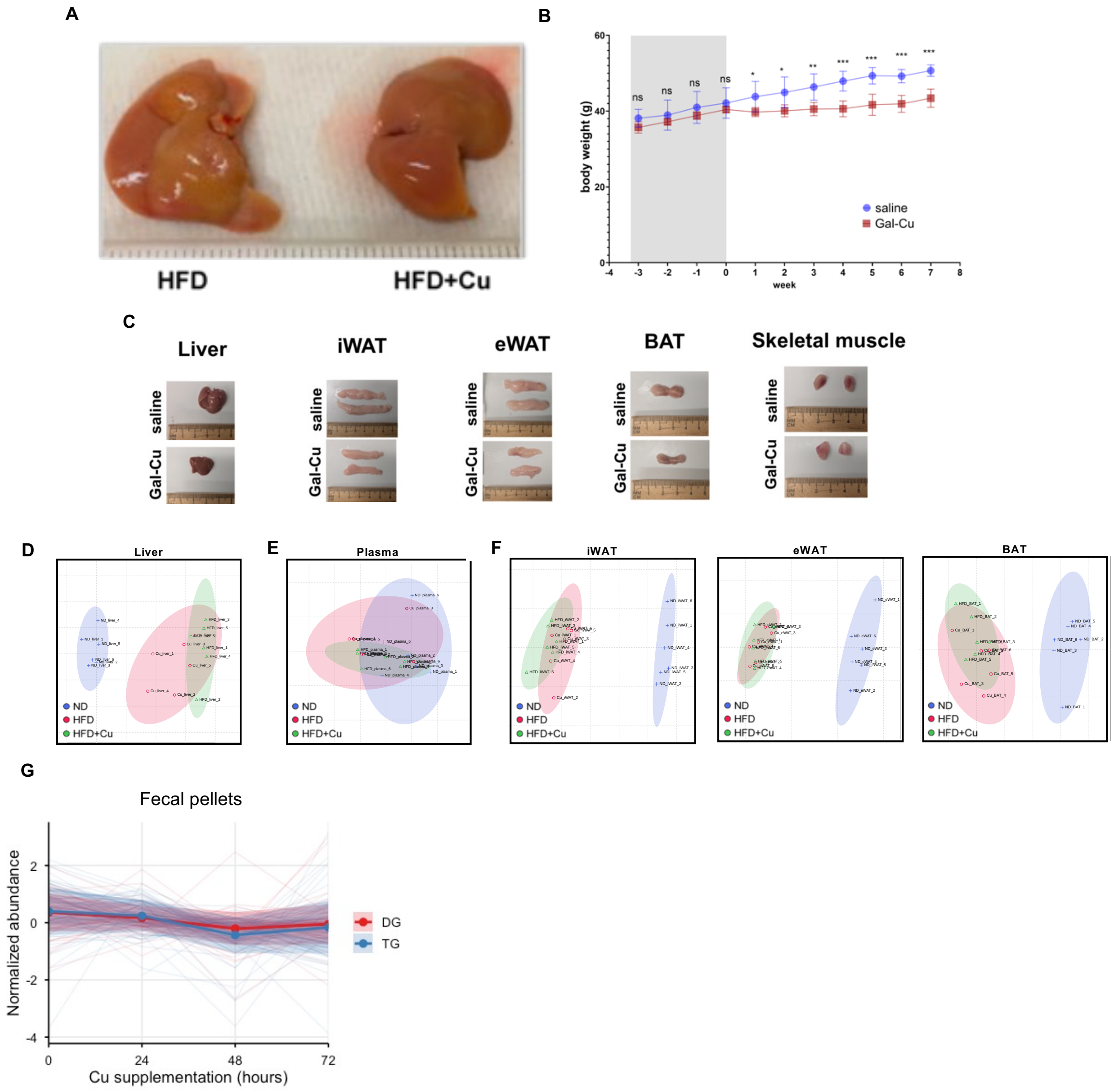


**Fig. S2: Copper supplementation in steatotic livers alter physiological phenotypes in mice.** **A.** Representative image of livers from mice fed with HFD for 2 weeks followed by Gal-Cu(gtsm) or DPBS for the following 10 weeks. **B-G.** Mice were fed normal diet or HFD for 12 weeks then normal diet for 8 weeks treatment with Gal-Cu(gtsm) or DPBS (n = 6). **B.** Mouse body weight measured before and during 8 weeks during copper supplementation for HFD-fed mice. Week 0 is the start of Gal-Cu(gtsm) treatment **C.** Representative images of tissues at the end point of therapeutic study of HFD-diet fed mice. **D-F.** PCA of di- and triglyceride lipid profiles in (D) liver, (E) plasma, and (F) adipose tissues. **G.** Normalized di- and triglyceride abundance in fecal pellets with 0, 24, and 48, and 72 hours of copper supplementation. Each light-colored line represents a lipid species. The solid line and shaded region represent mean ± SD of the DGs and TGs. All error bars represent mean ± SD. Anova; ns, not significant, $*p < 0.05$, $**p < 0.01$, $***p < 0.001$, $****p < 0.0001$ or exact numbers indicated.

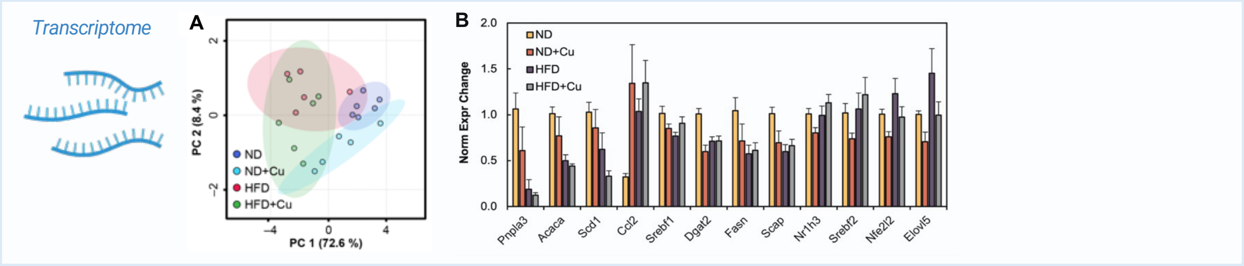


**Fig. S3: RNA transcripts of HFD and copper supplementation.**
**A-B.** Analysis of liver from mice that were fed with ND or HFD for 2 weeks followed by treatment with Gal-Cu(gtsm) or DPBS vehicle for the following 10 weeks (Fig. 2B, ND, HFD+Cu, n = 6; ND+Cu, HFD, n = 5 per group). **A.** PCA of mouse liver RT-qPCR transcripts. **B.** RT-qPCR of 12 MASLD-associated transcripts.

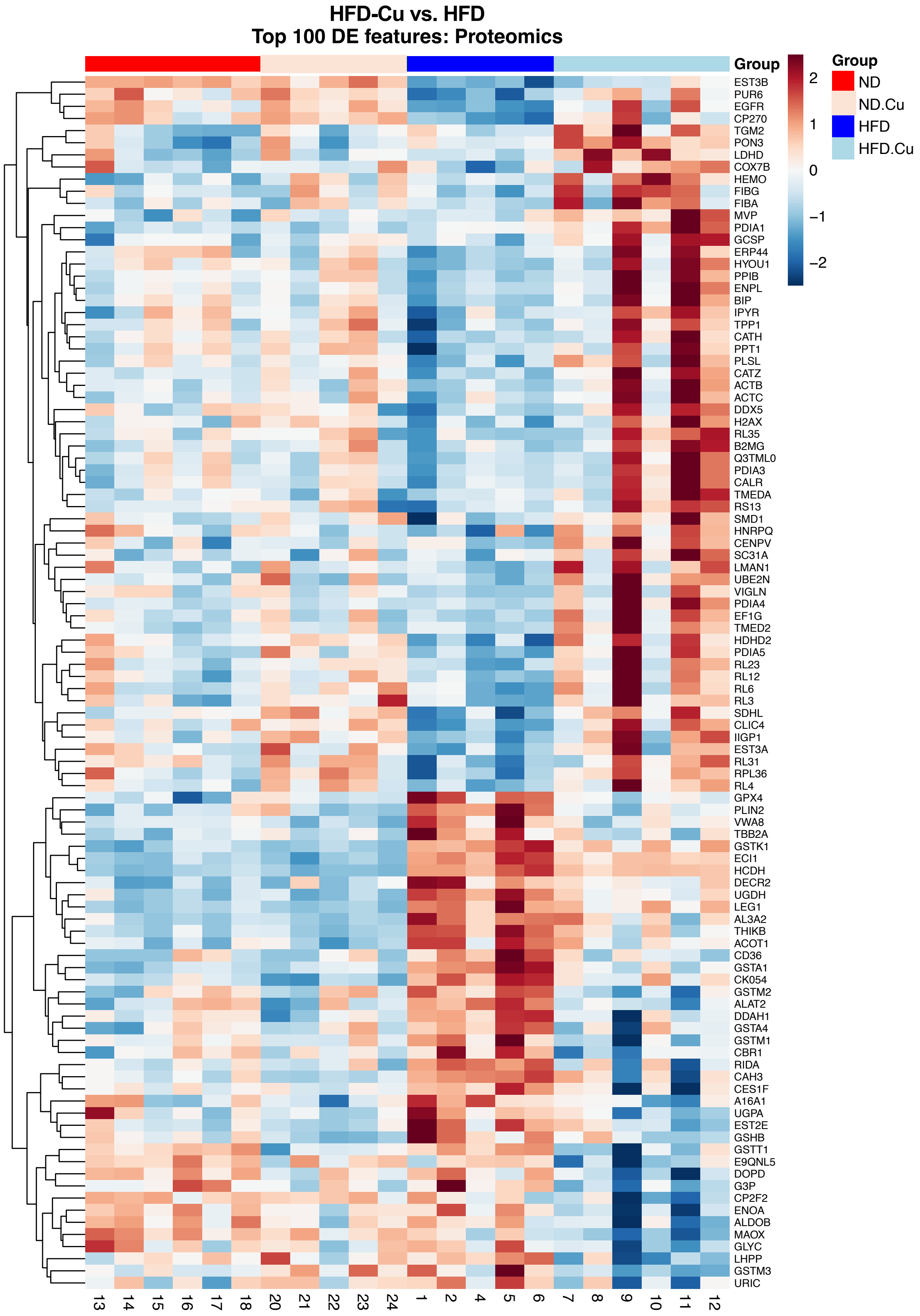


**Fig. S4: Heatmap of proteomics top 100 DEG comparing HFD+Cu vs. HFD.**

The liver proteome from mice fed with ND or HFD for 2 weeks followed by Gal-Cu(gtsm) or DPBS for the following 10 weeks (ND, HFD+Cu, n = 6; ND+Cu, HFD, n = 5) were analyzed. Heatmap shows top 100 proteins ordered by HFD+Cu vs. HFD. The color bar represents z-scores.

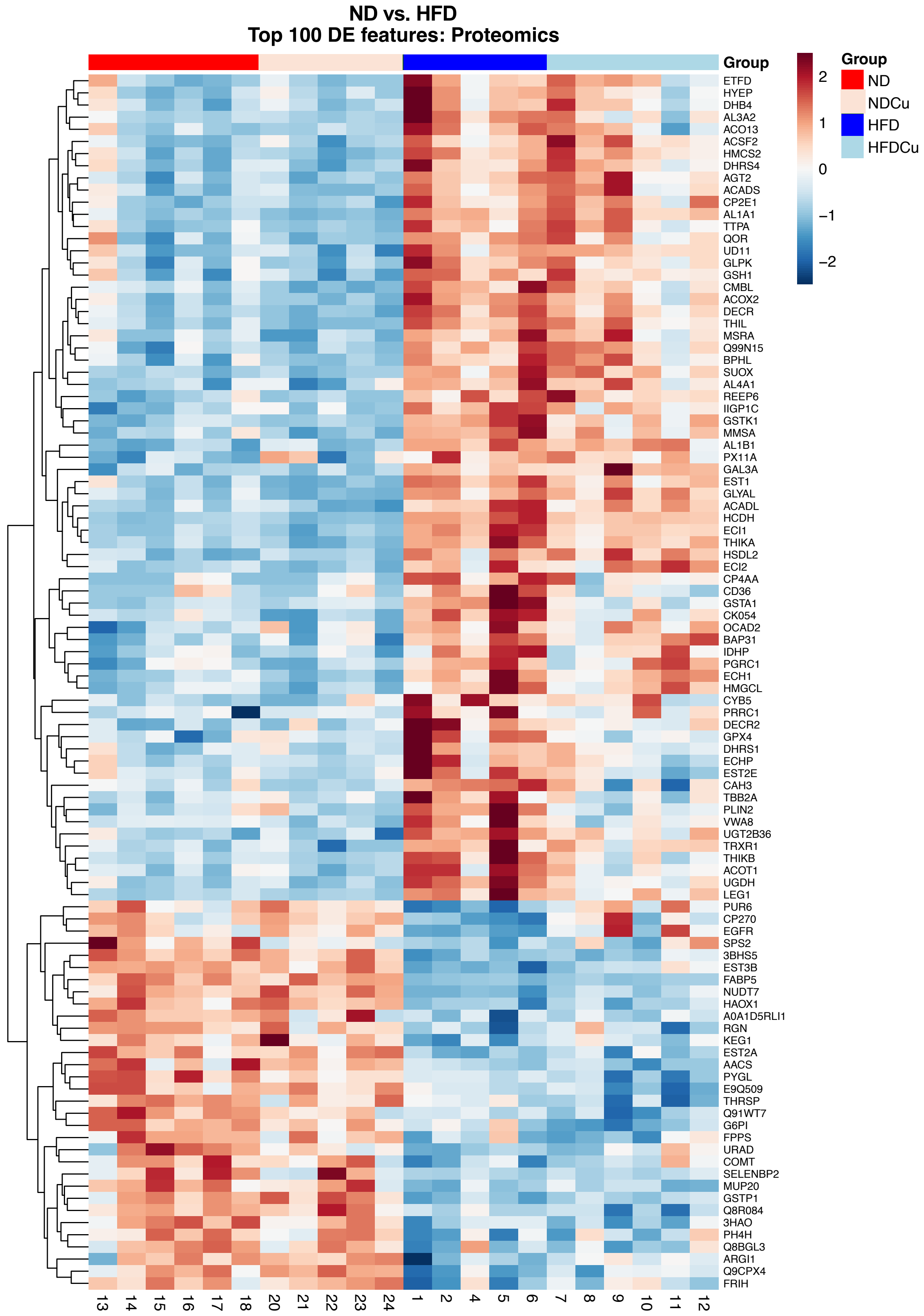


**Fig. S5: Heatmap of proteomics top 100 DEG comparing HFD vs. ND.**

The liver proteome from mice fed with ND or HFD for 2 weeks followed by Gal-Cu(gtsm) or DPBS for the following 10 weeks (ND, HFD+Cu, n = 6; ND+Cu, HFD, n = 5) were analyzed. Heatmap shows top 100 proteins ordered by ND vs. HFD. The color bar represents z-scores.

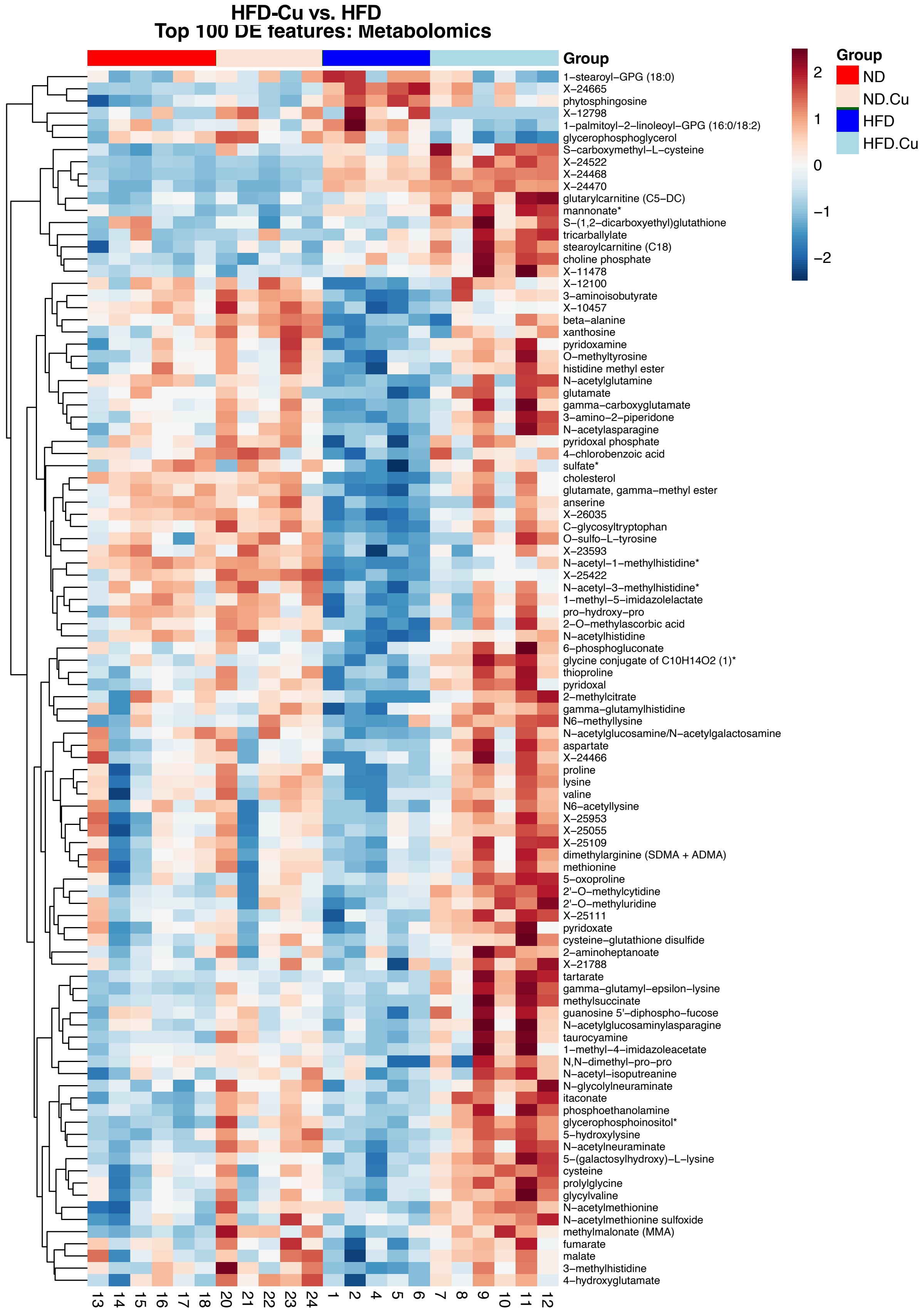
HFD-Cu vs. HFD
Top 100 DE features: Metabolomics
Group
2
1
0
−1
−2
Group
ND
ND.Cu
HFD
HFD.Cu
1−stearoyl−GPG (18:0)
X−24665
phytosphingosine
X−12798
1−palmitoyl−2−linoleoyl−GPG (16:0/18:2)
glycerophosphoglycerol
S−carboxymethyl−L−cysteine
X−24522
X−24468
X−24470
glutarylcarnitine (C5−DC)
mannonate*
S−(1,2−dicarboxyethyl)glutathione
tricarballylate
stearoylcarnitine (C18)
choline phosphate
X−11478
X−12100
3−aminoisobutyrate
X−10457
beta−alanine
xanthosine
pyridoxamine
O−methyltyrosine
histidine methyl ester
N−acetylglutamine
glutamate
gamma−carboxyglutamate
3−amino−2−piperidone
N−acetylasparagine
pyridoxal phosphate
4−chlorobenzoic acid
sulfate*
cholesterol
glutamate, gamma−methyl ester
anserine
X−26035
C−glycosyltryptophan
O−sulfo−L−tyrosine
X−23593
N−acetyl−1−methylhistidine*
X−25422
N−acetyl−3−methylhistidine*
1−methyl−5−imidazolelactate
pro−hydroxy−pro
2−O−methylascorbic acid
N−acetylhistidine
6−phosphogluconate
glycine conjugate of C10H14O2 (1)*
thioproline
pyridoxal
2−methylcitrate
gamma−glutamylhistidine
N6−methyllysine
N−acetylglucosamine/N−acetylgalactosamine
aspartate
X−24466
proline
lysine
valine
N6−acetyllysine
X−25953
X−25055
X−25109
dimethylarginine (SDMA + ADMA)
methionine
5−oxoproline
2'−O−methylcytidine
2'−O−methyluridine
X−25111
pyridoxate
cysteine−glutathione disulfide
2−aminoheptanoate
X−21788
tartarate
gamma−glutamyl−epsilon−lysine
methylsuccinate
guanosine 5'−diphospho−fucose
N−acetylglucosaminylasparagine
taurocyamine
1−methyl−4−imidazoleacetate
N,N−dimethyl−pro−pro
N−acetyl−isoputreanine
N−glycolylneuraminate
itaconate
phosphoethanolamine
glycerophosphoinositol*
5−hydroxylysine
N−acetylneuraminate
5−(galactosylhydroxy)−L−lysine
cysteine
prolylglycine
glycylvaline
N−acetylmethionine
N−acetylmethionine sulfoxide
methylmalonate (MMA)
fumarate
malate
3−methylhistidine
4−hydroxyglutamate
13 14 15 16 17 18 20 21 22 23 24 1 2 4 5 6 7 8 9 10 11 12

**Fig. S6: Heatmap of metabolomics top 100 DEG comparing HFD+Cu vs. HFD.** The liver metabolomes from mice fed with ND or HFD for 2 weeks followed by Gal-Cu(gtsm) or DPBS for the following 10 weeks (ND, HFD+Cu, n = 6; ND+Cu, HFD, n = 5) were analyzed. Heatmap shows top 100 metabolites ordered by HFD+Cu vs. HFD. The color bar represents z-scores.

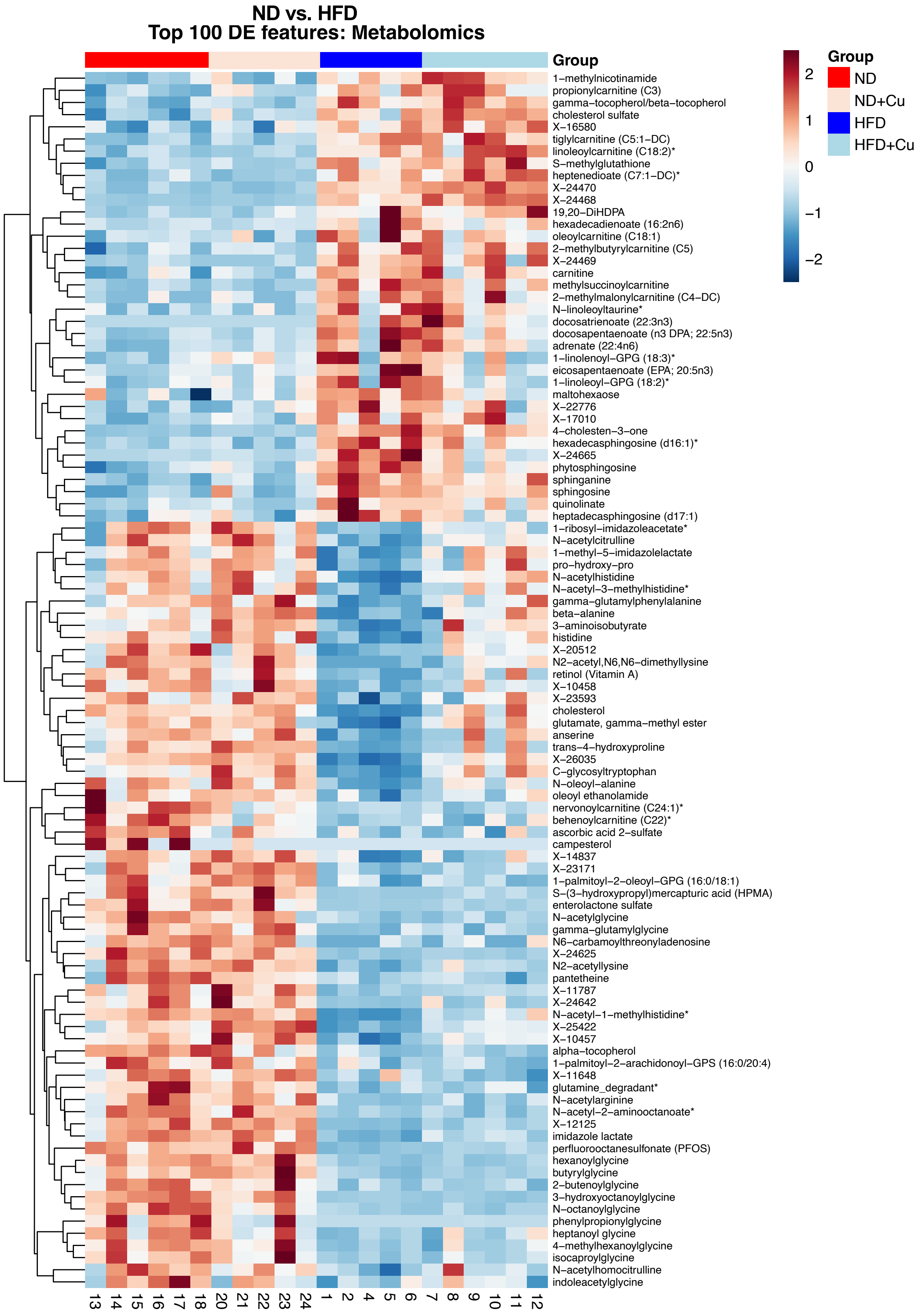
ND vs. HFD
Top 100 DE features: Metabolomics
Group
1−methylnicotinamide
propionylcarnitine (C3)
gamma−tocopherol/beta−tocopherol
cholesterol sulfate
X−16580
tiglylcarnitine (C5:1−DC)
linoleoylcarnitine (C18:2)*
S−methylglutathione
heptenedioate (C7:1−DC)*
X−24470
X−24468
19,20−DiHDPA
hexadecadienoate (16:2n6)
oleoylcarnitine (C18:1)
2−methylbutyrylcarnitine (C5)
X−24469
carnitine
methylsuccinoylcarnitine
2−methylmalonylcarnitine (C4−DC)
N−linoleoyltaurine*
docosatrienoate (22:3n3)
docosapentaenoate (n3 DPA; 22:5n3)
adrenate (22:4n6)
1−linolenoyl−GPG (18:3)*
eicosapentaenoate (EPA; 20:5n3)
1−linoleoyl−GPG (18:2)*
maltohexaose
X−22776
X−17010
4−cholesten−3−one
hexadecasphingosine (d16:1)*
X−24665
phytosphingosine
sphinganine
sphingosine
quinolinate
heptadecasphingosine (d17:1)
1−ribosyl−imidazoleacetate*
N−acetylcitrulline
1−methyl−5−imidazolelactate
pro−hydroxy−pro
N−acetylhistidine
N−acetyl−3−methylhistidine*
gamma−glutamylphenylalanine
beta−alanine
3−aminoisobutyrate
histidine
X−20512
N2−acetyl,N6,N6−dimethyllysine
retinol (Vitamin A)
X−10458
X−23593
cholesterol
glutamate, gamma−methyl ester
anserine
trans−4−hydroxyproline
X−26035
C−glycosyltryptophan
N−oleoyl−alanine
oleoyl ethanolamide
nervonoylcarnitine (C24:1)*
behenoylcarnitine (C22)*
ascorbic acid 2−sulfate
campesterol
X−14837
X−23171
1−palmitoyl−2−oleoyl−GPG (16:0/18:1)
S−(3−hydroxypropyl)mercapturic acid (HPMA)
enterolactone sulfate
N−acetylglycine
gamma−glutamylglycine
N6−carbamoylthreonyladenosine
X−24625
N2−acetyllysine
pantetheine
X−11787
X−24642
N−acetyl−1−methylhistidine*
X−25422
X−10457
alpha−tocopherol
1−palmitoyl−2−arachidonoyl−GPS (16:0/20:4)
X−11648
glutamine_degradant*
N−acetylarginine
N−acetyl−2−aminooctanoate*
X−12125
imidazole lactate
perfluorooctanesulfonate (PFOS)
hexanoylglycine
butyrylglycine
2−butenoylglycine
3−hydroxyoctanoylglycine
N−octanoylglycine
phenylpropionylglycine
heptanoyl glycine
4−methylhexanoylglycine
isocaproylglycine
N−acetylhomocitrulline
indoleacetylglycine
13 14 15 16 17 18 20 21 22 23 24 1 2 4 5 6 7 8 9 10 11 12
2
1
0
−1
−2
Group
ND
ND+Cu
HFD
HFD+Cu

1

**Fig. S7: Heatmap of metabolomics top 100 DEG comparing HFD vs. ND.**
The liver metabolomes from mice fed with ND or HFD for 2 weeks followed by Gal-Cu(gtsm) or DPBS for the following 10 weeks (ND, HFD+Cu, n = 6; ND+Cu, HFD, n = 5) were analyzed. Heatmap shows top 100 metabolites ordered by ND vs. HFD. The color bar represents z-scores.

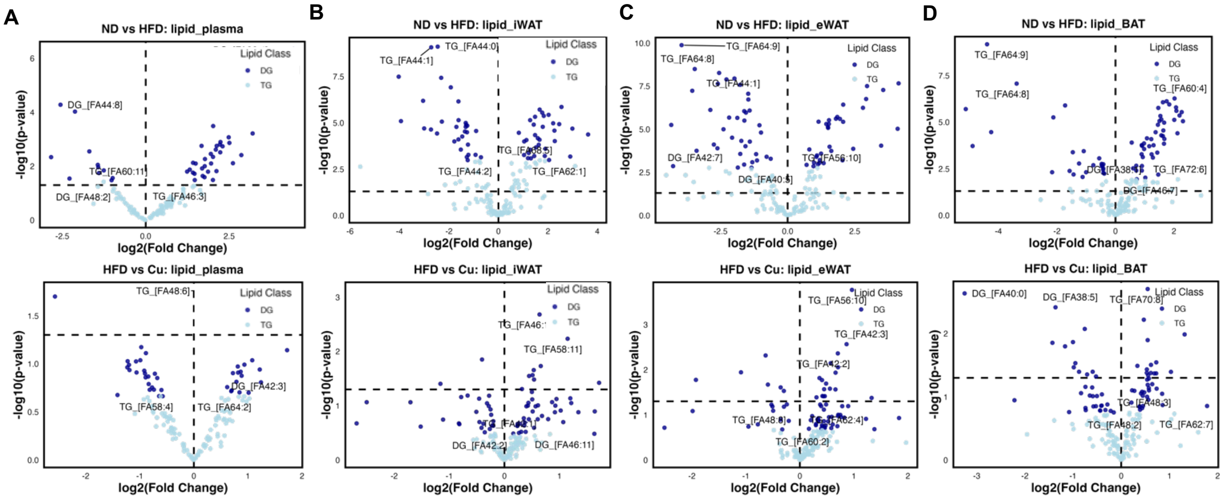


**Fig. S8: Lipidomics of liver-neighboring adipose tissues and plasma.**

**A-D.** The tissues from mice fed with ND or HFD for 12 weeks followed by Gal-Cu(gtsm) or DPBS for the following 8 weeks with ND were collected for analyzing lipidomics, and the volcano plots drawn with TG and DG classes. **A.** Volcano plots of ND vs. HFD and HFD vs. Cu plasma lipidomics. **B.** Volcano plots of ND vs. HFD and HFD vs. Cu iWAT lipidomics. **C.** Volcano plots of ND vs. HFD and HFD vs. Cu eWAT lipidomics. **D.** Volcano plots of ND vs. HFD and HFD vs. Cu BAT lipidomics.

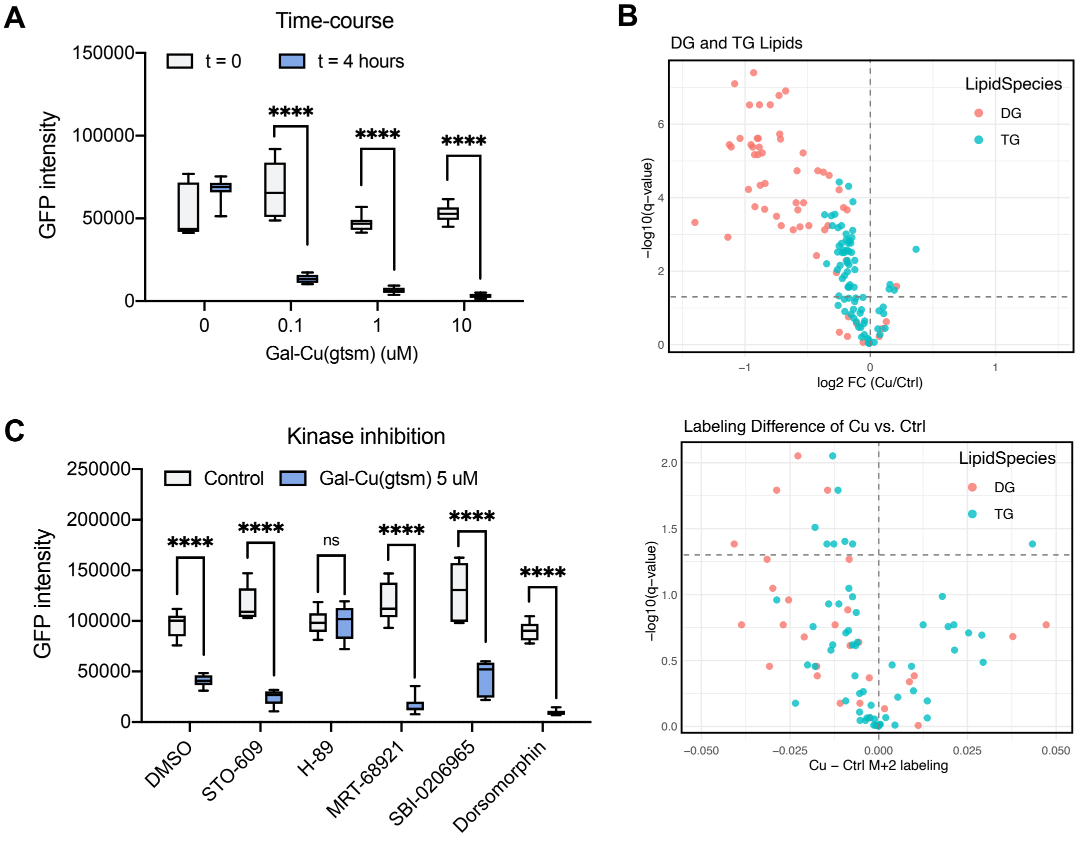


**Fig. S9: Copper effects metabolome and lipidome in hepatocytes.**
**A.** Live-cell imaging of PLIN2-GFP-Huh7 cells traced for 4 hours after indicated treatment. **B.** De novo lipogenesis of PLIN2-GFP-Huh7 cells. Cells were incubated with unlabeled acetate (top) or heavy-acetate (bottom) for 24 hours for labeling followed by Gal-Cu(gtsm) treatment for 4 hours and analyzed. **C.** Live-cell imaging of PLIN2-GFP-Huh7 cells stained with Hoechst 33342 (blue) and lysotracker (red) after treatment for 4 hours as indicated.

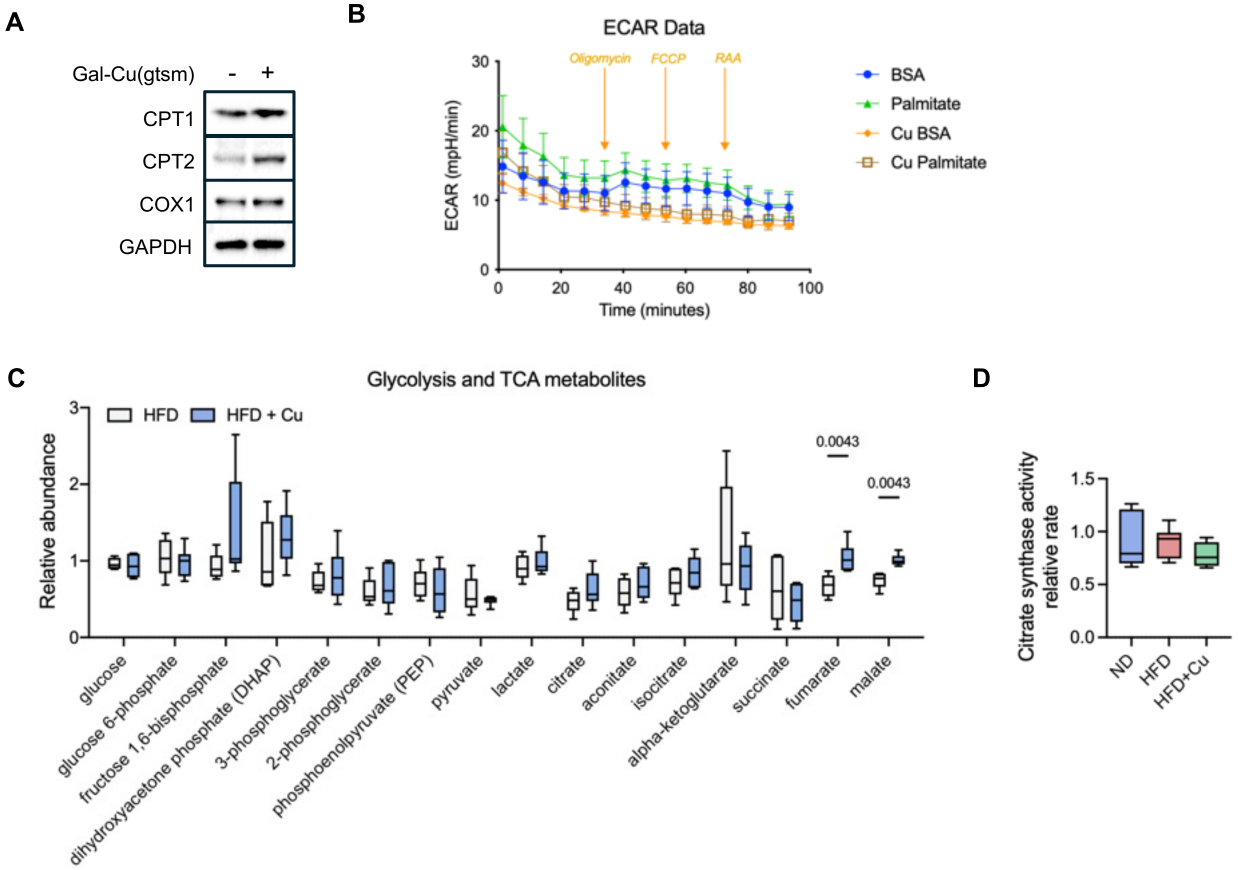


**Fig. S10: Copper effects metabolome and lipidome in hepatocytes.**

**A.** Immunoblotting of Huh7 cells treated with 1 mM of Gal-Cu(gtsm) or DMSO for 24 hours. **B.** ECAR measurement from seahorse assays on Huh7 cells after 4 hours of pretreatment of vehicle or Gal-Cu(gtsm). All error bars represent mean ± SD. **C.** The relative metabolite abundance and protein expression levels in liver from mice fed with ND or HFD for 2 weeks followed by Gal-Cu(gtsm) or DPBS for the following 10 weeks (ND, HFD+Cu, n = 6; ND+Cu, HFD, n = 5) measured. **D.** Relative enzyme activity measured in mitochondria suspension isolated from mouse livers from mice fed with ND or HFD for 2 weeks followed by Gal-Cu(gtsm) or DPBS for the following 10 weeks (ND, HFD+Cu, n = 6; ND+Cu, HFD, n = 5).
